# Integrative Spatial Single-cell Analysis with Graph-based Feature Learning

**DOI:** 10.1101/2020.08.12.248971

**Authors:** Junjie Zhu, Chiara Sabatti

## Abstract

We propose `GLISS`, a strategy to discover spatially-varying genes by integrating two data sources: (1) spatial gene expression data such as image-based fluorescence *in situ* hybridization techniques, and (2) dissociated whole-transcriptome single-cell RNA-sequencing (scRNA-seq) data. `GLISS` utilizes a graph-based association measure to select and link genes that are spatially-dependent in both data sources. `GLISS` can discover new spatial genes and recover cell locations in scRNA-seq data from landmark genes determined from SGE data. `GLISS` also offers a new dimension reduction technique to cluster the genes, while accounting for the inferred spatial structure of the cells. We demonstrate the utility of `GLISS` on simulated and real datasets, including datasets on the mouse olfactory bulb and breast cancer biopsies, and two spatial studies of the mammalian liver and intestine.

## 1 Introduction

Gene expression is often associated with the spatial structure of a tissue. The locations of the cells within the tissue can induce different states or expression gradients, even when we consider a single cell type [1, 2]. For instance, hepatocytes in the liver and enterocytes in the intestine are known to express certain genes in a spatially-dependent manner [3, 4] (Figure 1). To fully understand such phenomena, we need to comprehensively identify spatially varying genes.

**Figure 1:**
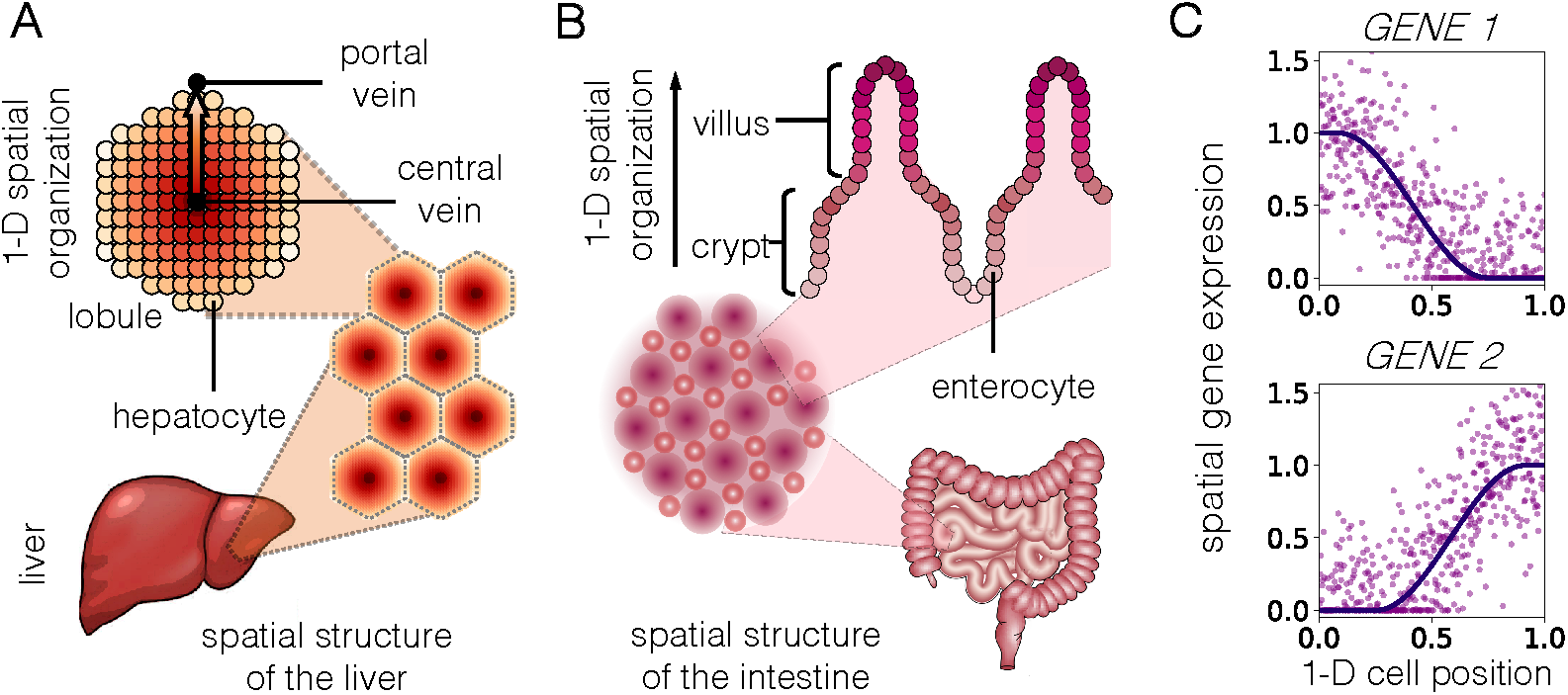
Examples of tissues with spatial gene expression studied in this paper. (A) Spatial structure of the liver. (B) Spatial structure of the intestine. (C) Two artificially generated gene expression profiles that vary spatially according to a 1-D cell position.

The advancement of single-cell RNA sequencing (scRNA-seq) technologies, which measure the whole-transcriptome at the single-cell resolution [5], have offered a new opportunity to analyze the roles of genes in distinct cells. Current scRNA-seq techniques dissociate individual cells into droplets or wells, resulting in the loss of spatial information [6, 7].

However, recent advances in spatial gene expression (SGE) profiling technologies have been able to measure gene expression at increasing resolution while preserving spatial information [8, 9]. For instance, this is the case with image-based fluorescence *in situ* hybridization (FISH) techniques, such as seqFISH [10, 11] and MERFISH [8, 12]. These technologies have been used to analyze the embryo [13], the brain [14], or even subregional biopsies from cancer samples [15]. Current SGE technologies have been restricted in two ways: first, some technologies can only measure a small number of genes on the same spatial coordinates due to a limited number of multiplexed channels; second, the throughput of measured cells is often constrained by the sampled region and image resolution. The lower efficiency of SGE measurements remains a shortcoming when compared to dissociated scRNA-seq, which measures the whole-transcriptome at higher cell throughput and lower cost. Because SGE techniques do not measure all possible genes, spatially varying genes that we aim to discover could be already excluded from the data analysis. Moreover, statistical methods can still fail to detect spatially varying genes that are measured in SGE data when the sample size is too small.

Because our goal is to comprehensively mine spatial gene expression patterns, we need expression of all genes over a large sample size, while maintaining the spatial information. This can be done by combining the information across scRNA-seq and SGE data. In fact, the absence of positional information in scRNA-seq has been compensated by linking SGE data of the same tissue in some recent studies.

In a recent study [3], hepatocytes in the mammalian liver were analyzed to discover spatially varying genes with central-portal biases (Figure 1A). To combine the two data sources, the analysis workflow: (1) divided the central-portal axis measured by the SGE data into nine zones, (2) identified six landmark genes that are differentially expressed across the zones, (3) built a classifier to map each cell from the scRNA-seq data to each of the nine zones based on the six landmark genes, and (4) assessed if other genes were differentially expressed across the central-portal axis based on the predicted cell locations. The study proposed and validated novel spatially varying genes, and, in particular, ones whose variation in expression was non-monotonic with respect to the inferred positions along the central-portal axis.

This selection procedure required manual curation and thus highly specific to the study. For instance, they selected non-monotonic genes based on the arbitrary criteria that the expression peaks between zones 3 and 7, and that this peak value needs to be 10% higher than the maximal value between layers 1 and 9. Such manual curation can prohibit the reproducibility and scalability of this analysis.

Another limitation of the previously mentioned study and others [13, 4] is that the previous analysis discretized and partitioned of the cells based on spatial positions. Indeed, such partitioning is often highly data-dependent and not necessarily replicable in a new study because the number of groups or cross-group cell type proportions can differ. Besides, the number of positional groups is a hyperparameter, to which the gene selection process has high sensitivity. Determining its value is complex: if the number of groups is too small, then the spatial structure within each group can potentially be obscured; if the number of groups is too large, then the number of comparisons across the them becomes too large to guarantee a statistically powerful analysis.

### 1.1 Our Contribution

We introduce `GLISS` (**G**raph **L**aplacian-based **I**ntegrative **S**ingle-cell **S**patial Analysis), a graph-based feature selection and interpretation framework that allows us to

- identify spatially varying genes genes from SGE data as landmark genes;
- leverage these landmark genes to discover new spatially varying genes in scRNA-seq data;
- infer latent spatial structures among the cells in scRNA-seq data;
- reduce the dimension of the gene observations and cluster genes based on their spatial patterns.

We treat the cell position as a continuous variable for the following reasons: (i) it captures the continuous nature of the expression gradients more faithfully, (ii) it is more generalizable to other studies because it removes a data-dependent preprocessing step, and (iii) it eliminates the need of choosing the discretization hyperparameter.

`GLISS` includes a principled and generalizable spatially varying gene selection process: we use a graph-based correlation measure that automatically selects both monotonic and non-monotonic associations between multivariate spatial variables and gene expression information. The procedure is model-free: we do not make distributional assumptions about the data generating process for either SGE or scRNA-seq data. Further, our non-parametric statistical procedures are empirically powerful with false discovery guarantees, which is crucial for generalizability and reproducibility.

### 1.2 Related Work

Recently, an increasing number of works have focused on data integration across multiple single-cell experiments as more scRNA-seq datasets have become available [16, 17, 18, 19]. Although most of these works are primarily designed for batch effect correction [19] (when concatenating different scRNA-seq datasets), some of them have extended their methodology to integrate SGE and scRNA-seq data. Seurat 3.0 [17] proposed an integrative analysis workflow that relies on subsets of cells, also known as ‘anchors’, which can be linked across datasets. ssGPLVM [18] proposed an alternative probabilistic approach to perform data integration while allowing for more complex covariates that capture the batch difference between the SGE and scRNA-seq data. Both Seurat and ssGPLVM can be seen as approaches to augment the sample size in the SGE data by leveraging the high-throughput scRNA-seq data, because they map the cells in scRNA-seq data to the explicit coordinates measured in SGE data.

Rather than linking the *cells* across two data sources, `GLISS` links the data through a common set of *genes* that contain spatial information. This is done by identifying spatially varying genes from SGE data and then leveraging them in scRNA-seq data as landmark genes. To align cells across datasets, one needs multiple distinct cell types, whereas `GLISS` focuses on detecting fine-grained spatially varying genes within a single cell type. In fact, `GLISS` does not require the samples in the SGE data to correspond to individual cells. For instance, the SGE samples can correspond to spatial bulk RNA-seq measurements, i.e., aggregated gene expression of cells within a region. Secondly, cell alignment strategies heavily rely on which genes are available in SGE data, potentially ignoring the information contained in the additional genes measured in scRNA-seq data. `GLISS` takes into account all genes present in each data source, which allows us to also detect spatially varying genes in scRNA-seq data. Further, `GLISS` relies on a non-parametric spatial model that is robust to model misspecification and allows to define rigorous statistical tests.

The workflow of `GLISS` includes multiple sub-components including (1) detecting spatially varying genes in SGE data alone, and (2) inferring latent trajectories among cells. There are two lines of work that relate to these specific sub-components.

Recently, systematic methods been introduced for spatially varying gene detection from SGE data alone: `SpatialDE` [20] adopts a linear mixed effects model from statistical genetics to decompose the covariance of the observed gene expression into the spatial and non-spatial components, and then it determines the significance of the spatial variability based on a likelihood test; `trendsceek` [21] models the spatial expression of a gene via a marked point process from geostatistics, and then relies on four non-parametric summary statistics to determine the spatial significance of a gene compared to a null empirical model; and more recently, `scGCO` [22] was introduced to address the problem of scalability. Interestingly, `scGCO` differs from the previous approaches as it uses a graph-based formulation constructing a Voronoi partition of the image in 2-D. These methods leverage explicit positional information, and therefore, cannot apply to scRNA-seq data when the spatial coordinates are absent. One could apply the existing SGE methods to detect landmark genes in the workflow of `GLISS`; in addition, we offer a common graph-based approach that can be consistently applied to both SGE and scRNA-seq data. We will compare the performance of our approach with these SGE methods on simulated and real datasets.

The other line of related work focus on trajectory inference. The concept of ‘pseudotime’ was introduced to describe the temporal relationship of the cells captured in one snapshot in dissociated scRNA-seq data [23]. Initial proposals to infer such cell orderings have then been followed by the development of over 70 trajectory inference methods [24, 25]. The inference of a onedimensional temporal relationship shares certain similarities with that of spatial relationships along a one-dimensional latent axis, which is the focus of our work. In fact, the spatial structures can sometimes coincide with the temporal relationships among cells (i.e., those that differentiate along a spatial axis). The existing trajectory inference methods are tailored to detect a strong global effect (e.g., differentiation signals) shared across the majority of the genes, whereas `GLISS` selectively targets a subset of genes that contain spatial information.

## 2 Method

The common inputs for `GLISS` are: (1) the spatial gene expression (SGE) matrix with matched spatial coordinates (Figure 2A) and (2) the scRNA-seq gene expression matrix (Figure 2B). Each row of the matrices correspond to a cell but the samples across the two data sources are not matched: the rows in the SGE matrix do not correspond to any row in the scRNA-seq matrix, and vice versa. The genes (columns) in both matrices may not be identical, especially because SGE measurements can be limited to a small set of genes. However, we require the genes in both matrices to overlap to some extent, since `GLISS` leverages some of the spatially varying (SV) genes in SGE as LM genes to analyze the scRNA-seq data.

**Figure 2:**
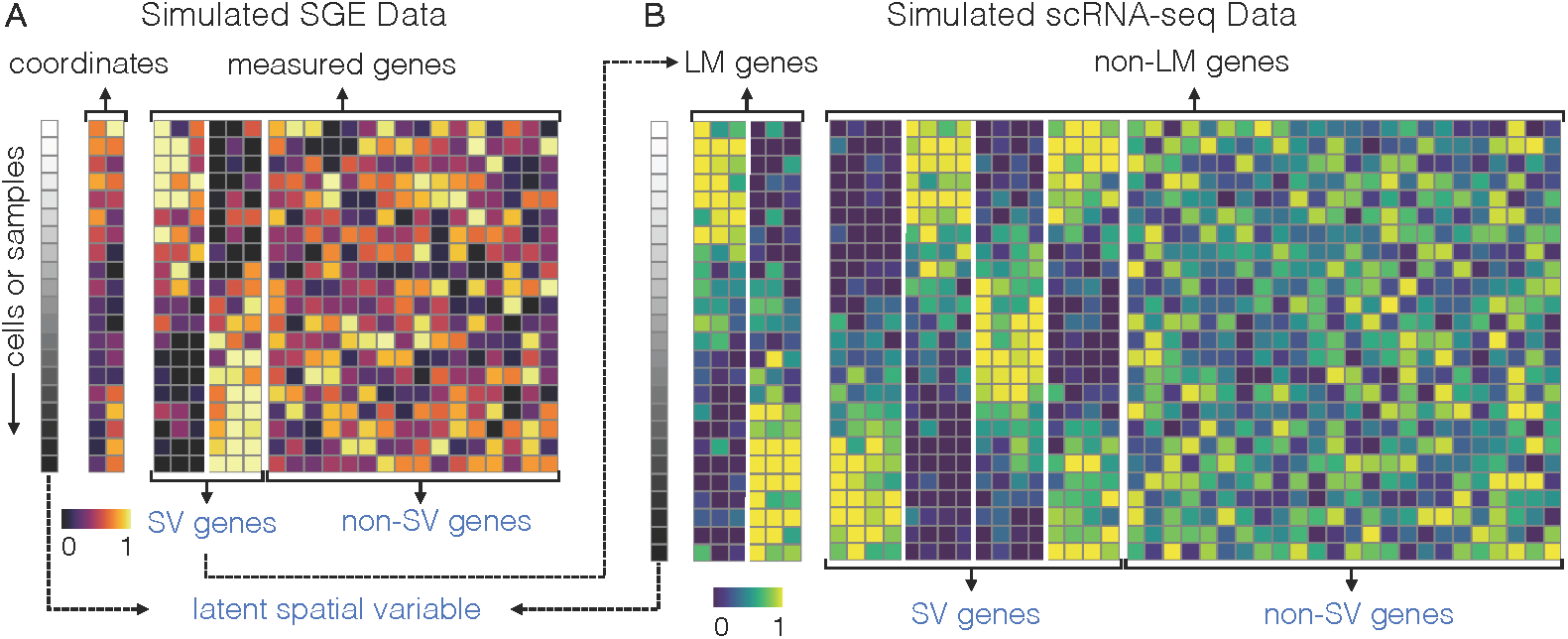
Illustration of observed (black) and unobserved (blue) components of SGE and scRNA-seq data. (A) SGE data linked with observed spatial coordinates. (B) scRNA-seq data divided by landmark (LM) genes and non-LM genes. The LM genes are determined for the scRNA-seq are the spatially varying (SV) genes identified in the SGE analysis.

Under certain scenarios, such as spatially-captured bulk RNA-seq, each row in SGE can also represent a collection of cells, whose average expression is measured over an indexed spatial region. Throughout this paper, we will still refer to these samples as “cells” for the SGE analysis, as `GLISS` detects SV genes the same way in all the scenarios. Note that our workflow can also accommodate the scenario where SGE data is unavailable but a set of SV genes are already provided as input LM genes for the scRNA-seq analysis.

The remainder of this section is structured as follows: first, we provide a brief overview of `GLISS` given the two data sources (Section 2.1); then, we introduce our integrative model and technical assumptions about the data (Section 2.2); finally, we elaborate on details about specific components of `GLISS` and discuss several theoretical properties (Section 2.3).

### 2.1 Overview of GLISS

Figure 3 illustrates the full workflow of `GLISS` given these inputs. The workflow consists of the following components:

① For the input SGE data, `GLISS` constructs a neighborhood graph from the spatial coordinates: each node represents a cell and two nodes share an edge if they are spatially close to each other ^1^. Then `GLISS` applies a graph-based feature selection procedure to determine SV genes that are dependent on the graph structure, and declare them as LM genes. `GLISS` can also directly take in as input a set of user-defined LM genes and skip this step of the analysis, meaning researchers can rely on existing literature to determine LM genes from other SGE data analyses.
② For the input scRNA-seq data, the cells could be selected from known markers or isolated populations determined from a clustering procedure. As we view this procedure as a preprocessing step (along with quality control) prior to running `GLISS`, we use subroutines from well-established scRNA-seq pipelines such as Seurat [28] or scanpy [29].
③ Given the curated scRNA-seq data from ②, `GLISS` constructs a cell-to-cell similarity graph based on the expression of the LM genes identified from ①, such that two nodes representing cells share an edge if they are close in the LM gene expression space. This similarity graph serves as a surrogate of the spatial similarity since cells in the scRNA-seq data do not contain spatial coordinates. Similar to ①, `GLISS` uses the constructed graph to select SV genes among the non-LM genes.
④ `GLISS` provides the option to infer a latent graph embedding for the cells based on both the newly selected SV genes and the LM genes. Using the graph spectrum, `GLISS` allows one to predefine an arbitrary dimension size to generate the latent embedding. In the hepatocytes or enterocytes examples (Figure 1), it can be useful to re-parametrize and order each cell with a 1-D embedding.
⑤ `GLISS` allows one to visualize and cluster genes in scRNA-seq, while accounting for their spatial dependency. It relies on spline models to fit each gene’s expression on the inferred latent embedding to learn their explicit spatial dependency. Then, `GLISS` leverages the fitted coefficients to reduce the dimensionality of each gene from the number of cells to the number of coefficients, so that similar SV genes can be clustered together.

**Figure 3:**
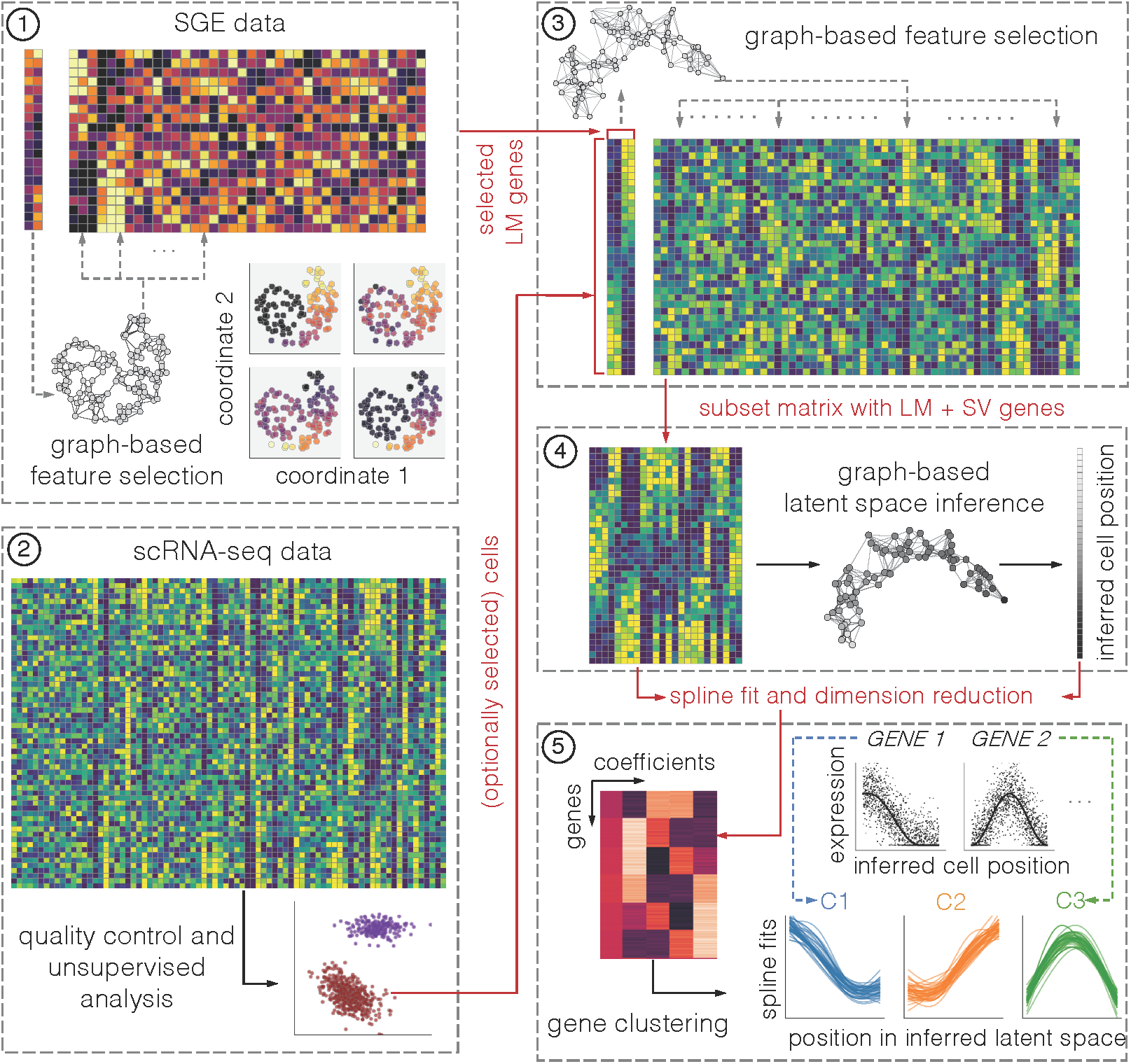
Overview of the workflow of `GLISS`. ① It selects SV genes in SGE data as LM genes using graph-based feature selection. ② It relies on quality control and unsupervised learning methods to determine cells of interest in the scRNA-seq data. ③ It determines new SV genes in scRNA-seq data also using graph-based feature selection. ④ It applies a graph-based approach to reconstruct latent cell positions using the selected SV genes. ⑤ After fitting each gene against the latent cell positions, it provides a low-dimensional representation for each gene based on the fitted coefficients and performs clustering in the coefficient space.

### 2.2 Integration Model and Assumptions

#### 2.2.1 Spatial Gene Expression Model

Given SGE data that measures the expression of *n* cells, let 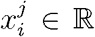 and *s*_*i*_ ∈ ℝ^*ℓ*^ be the expression of gene *j* and the spatial coordinate of cell i, respectively^2^. The spatial coordinates are often measured in 2-D, i.e., *ℓ* = 2, but can also have other number of dimensions (Figure 2A). Here, we assume a nonparametric spatial model for each gene by

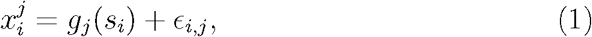

where function *g*_*j*_ captures the spatial dependency of gene *j* on the *ℓ*-dimensional coordinates, and the error term *ϵ*_*i,j*_captures non-spatial variability, such as technical noise in measuring the expression. As part of our nonparametric model, we do not assume we know the form of *g*_*j*_ or the distribution of *ϵ*_*i,j*_.

We formally assume that, given the ordered spatial coordinates *s*_1_, *s*_2_,…, *s*_*n*_, the random variables *ϵ*_1;*j*_, *ϵ*_2,*j*_, …,*ϵ*_*n,j*_ are *exchangeable*, i.e., their joint distribution of *ϵ*_1,*j*_, *ϵ*_2,*j*_, …,*ϵ*_*n,j*_ does not change if we swap any two of them. In other words, the error terms *ϵ*_1,*j*_, *ϵ*_2,*j*_, …, *ϵ*_*n,j*_ capture the common source of variation in the expression that is not associated with the spatial location. The model imposes the assumption that systematic variability in the expression across different cells is only driven by the spatial locations. Therefore, we can interpret the model in the context of a single cell type which exhibits spatial expression changes, or multiple cell types whose organization corresponds exactly to their spatial locations.

For the scRNA-seq data measured on *N* different cells, we assume a similar spatial model for 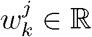 (e.g., the expression of gene *j* in cell *k):*

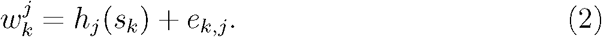

Notice that we alter the cell index in the scRNA-seq model to remember that the cells cannot be simultaneously measured by both SGE and scRNA-seq technologies. We replaced the error term *ϵ*_*i,j*_ with *e*_*k,j*_ because the noise characteristics differ between the two data sources. For instance, SGE data obtained by FISH can be corrupted by probe-specific noise [30], whereas droplet-based scRNA-seq data exhibit significant missing observations (known as ‘dropouts’) due to transcript capture efficiency [31]. Here, we also assume that given *s*_1_, *s*_2_,…, *s*_*N*_, the random variables *e*_1,*j*_, *e*_2,*j*_, …, *e*_*N,j*_ are exchangeable.

##### True underlying SV and non-SV genes

For the SGE and scRNA-seq data, we say that a gene is a non-SV gene if *g*_*j*_ (or *hj*) is a constant, and we say that the gene is an SV gene otherwise. We assume that there are true underlying SV genes and non-SV genes that are unknown a priori in both data sources (Figure 2). We also refer to the true underlying SV genes as *non-null features* and the non-SV genes as *null features*.

##### Linking datasets with LM genes

We assume that the two data sources share an underlying set of SV genes^3^, such that if gene *j* is in this set, it satisfies the following property: *g*_*j*_ is not a constant implies that hj is not a constant. This assumption has frequently been implicitly used to define underlying LM genes within the same biological system of interest [3, 4]: if the SGE data strongly suggests a gene to be spatially varying, we would also expect it to also be spatially varying when measured with scRNA-seq technology. Without this assumption, integrative analysis would not meaningful because neither the cells nor the genes from both data sources would have anything in common. To avoid ambiguity, we refer to the SV genes that are inferred from the SGE data (or validated in other studies) as LM genes when we consider them in the context of scRNA-seq data.

#### 2.2.2 Statistical Hypotheses for SV Gene Detection

SV gene detection is used in two steps: (1) identifying LM genes from SGE data, and (2) determining new SV genes from scRNA-seq data based on the selected LM genes from SGE analysis (Figure 3). Each time we determine if a gene is a null feature or not, we are investigating if its expression is spatially varying or not. However, because the spatial coordinates are available in SGE but not in scRNA-seq data, the null hypotheses for two data sources need to be formulated differently. Within in each data type, we will overload the notation *j* to index a gene and denote its null hypothesis with 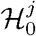.

##### Null hypothesis for SGE data

The measured expression level of gene *j* on *n* cells in SGE data comes with a matching set of spatial spatial coordinates, i.e., 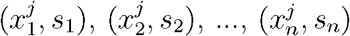. We state the null hypothesis by requiring that the joint distribution of 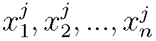 would not differ if their order is altered while fixing the spatial coordinates *s*_1_, *s*_2_,…, *s*_*n*_. In other words,

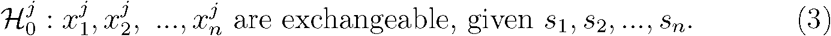

The SGE null hypothesis is stated to be model-free, but we can also interpret gene *j* as a null feature in the context of the data generative model.

###### Proposition 2.1.

*The null hypothesis* 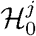 *in* *Eq*. (3) *is true if and only if g*_*i*_ *is a constant under the model in* *Eq*. (1).

*Proof*. If *g*_*i*_ equals a fixed constant *C* under the model, then 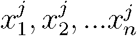 will be exchangeable because the exchangablity of *ϵ*_1,*j*_, *ϵ*_2,*j*_,… *ϵ*_*n,j*_ still holds after being shifted by a constant *C*; on the other hand, if *g*_*j*_ were not constant, we would identify two spatial locations *s*_*i*_ and *s*_*k*_ such that the marginal distributions of 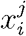 and 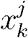 are not identical, which would imply that 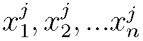 are not exchangeable. □

##### Null hypothesis for scRNA-seq data

In scRNA-seq data, we do not observe the spatial coordinates, but we assume we have obtained a set of LM genes from the SGE analysis. Let 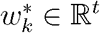 denote a length-t vector that include expression values of a set of *t* LM genes in cell *k*, and let *N* be the number of cells. The null hypothesis related to the non-LM gene *j* is stated as

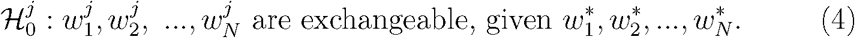

In other words, the expression of gene *j* is not associated with the expression of the LM genes under the null hypothesis. Note that, if we knew the unobserved spatial positions of each cell, i.e., *s*_1_, *s*_2_,…, *s*_*N*_, we would still directly test a null hypothesis in Eq. (3):

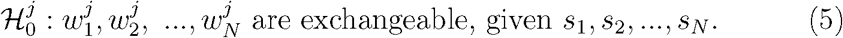

The relation between Eq. (4) and Eq. (5) can be understood in light of Eq. (2) with some additional assumptions below.

###### Proposition 2.2.

*Assuming the model in* *Eq*. (2) *and let* 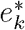 *be the length-t vector consisting of each error values of the t LM genes. If e*_1,*j*_,…, *e*_*N,j*_ *are exchangeable given* 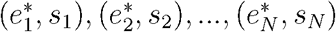, *the null hypothesis in* *Eq*. (5) *implies the null hypothesis in* *Eq*. (4).

*Proof*. If *e*_1,*j*_,…, *e*_*N,j*_ are exchangeable given 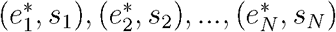, then *e*_1,*j*_,…, *e*_*N,j*_ are exchangeable given 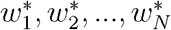 due to the model in Eq. (2). Further, if we shift each element in *e*_1,*j*_,…, *e*_*N,j*_ by a constant *C*, they will still remain exchangeable. According to Proposition 2.1, the null hypothesis in Eq. (5) implies that we have 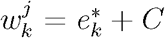 for each k and for a fixed *C*. Hence, the variables 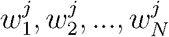 are exchangeable given 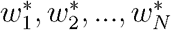. □

The exchangability assumption in Proposition 2.2 indicates that the measurement noise for gene *j* is unassociated with the noise of each of the LM genes. This is a strong assumption that we do not expect to be satisfied for every gene in real data analysis. For such genes whose noise is associated with the LM genes, the testable null hypothesis in Eq. (4) is no longer able to distinguish them from those that are associated with the LM genes due to the spatial structure. We will illustrate later via simulations (Sections 2.3.3 and 3.3) how our workflow can still be used to empirically analyze such data.

##### Selection among multiple hypotheses

Because we are considering multiple hypotheses (one per gene), we focus on two metrics to assess the feature selection quality: the statistical power and the False Discovery Proportion (FDP). We denote the sets of true null and non-null features as 𝒱^0^ and 𝒱^1^, respectively. Given ℛ, the set of the features selected from the data by a method, the (empirical) power and the FDP are defined as:

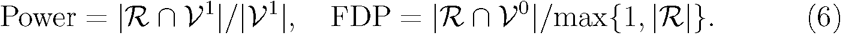

While we desire high power when selecting the non-null features, we control the proportion of false discoveries among the selected features. In particular, we aim to control the False Discovery Rate (FDR), defined as FDR = 𝔼[FDP], i.e., the average FDP over multiple realizations of the data.

#### 2.2.3 Latent Space Modeling Assumptions

The previous models and assumptions are sufficient to interpret the result of SV gene detection in SGE and scRNA-seq data. If one uses `GLISS` to reconstruct a latent structure from the scRNA-seq data or to cluster the SV genes according to this structure, we underline some additional assumptions that lead to meaningful interpretation of the results.

Instead of taking the absolute coordinates within the system to model a gene’s variability, we update Eq. (2) with a latent spatial factor (Figure 3B) such that spatial model becomes:

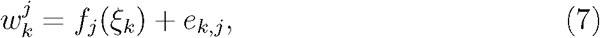

where *ξ*_*k*_ ∈ ℝ^*q*^ is a q-dimensional latent spatial variable so that we can substitute *h*_*j*_ (s_*k*_) with *f*_*j*_ (*ξ*_*k*_). Under this altered model, it still holds that gene *j* as a null feature if and only if *f*_*j*_ is a constant function, due to Proposition 2.1. Subsequently, the null hypothesis in Eq. (4) can still be re-interpreted as the necessary condition for 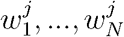 to be exchangeable given *ξ*_1_,…, *ξ*_*N*_, following the similar construction in Proposition 2.2.

##### Dimensionality of the latent space

The number of latent dimensions *q* is often user-defined, but in spatial analysis applications it is often sufficient to consider *q* = 1 or 2 since the spatial variable *s*_*k*_ is at most 3-dimensional. If *q* = 1, then the values of *ξ*_1_,…, *ξ*_*N*_ would order the cells along a 1-D axis, which is relevant to the hepatocyte and enterocyte applications (Figure 1).

##### Continuity assumptions

First, because we want to preserve the continuous nature of the spatial positions, we assume that each of *ξ*_1_,…, *ξ*_*N*_ is sampled from a continuous distribution. Second, to recover the latent spatial structure from the gene expression gradients, we need to assume that each SV gene expression varies continuously with the latent structure of interest, which is often the case for fine-grained tissue samples. More specifically, if gene *j* is an SV gene, the function fj is continuous at any point *ξ*_*k*_: for any neighborhood *N*_1_*(f*_*j*_(*ξ*_*k*_)), there exists a neighborhood *N*_2_*(ξ*_*k*_) such that *f*_*j*_*(z)* ∈ *N*_1_*(f (ξ*_*k*_)) whenever *z* ∈ *N*_2_*(ξ*_*k*_). Because we will use a graph-based model to infer *ξ*_*k*_, it is convenient to consider this (neighborhood) continuity definition, which we interpret as: if two cells that are close in the latent space, then their expression values of a true SV gene should also be similar.

### 2.3 Details on Specific Procedures

Here we elaborate on the details of several subroutines of Figure 3: the graph-based feature selection procedure (Steps ① and ③), graph-based latent space inference (Step ④, and gene dimension reduction and clustering (Step ⑤).

#### 2.3.1 Graph-based Feature Selection

One novely of `GLISS` is that it constructs graphs from expression data to capture cell-to-cell similarities, and then relies on the spectral properties of the graphs to perform feature selection (as well as latent space inference). Spectral techniques have been popular in spectral clustering [26] due to their sensitivity to local structures, and we reasoned that in our application, a lot of spatial gene variability can indeed be highly localized within a measured region.

To select SV genes in both SGE and scRNA-seq data, `GLISS` relies on the same graph-based non-parametric approach (Algorithm 1), derived from the Laplacian Score (LS) [27], permutation testing, and the Benjamini-Hochberg (BH) procedure [32]. Below, we introduce the method with reference to SGE data, but the same applies to the scRNA-seq data.

##### Algorithm 1: Feature Selection with GLISS

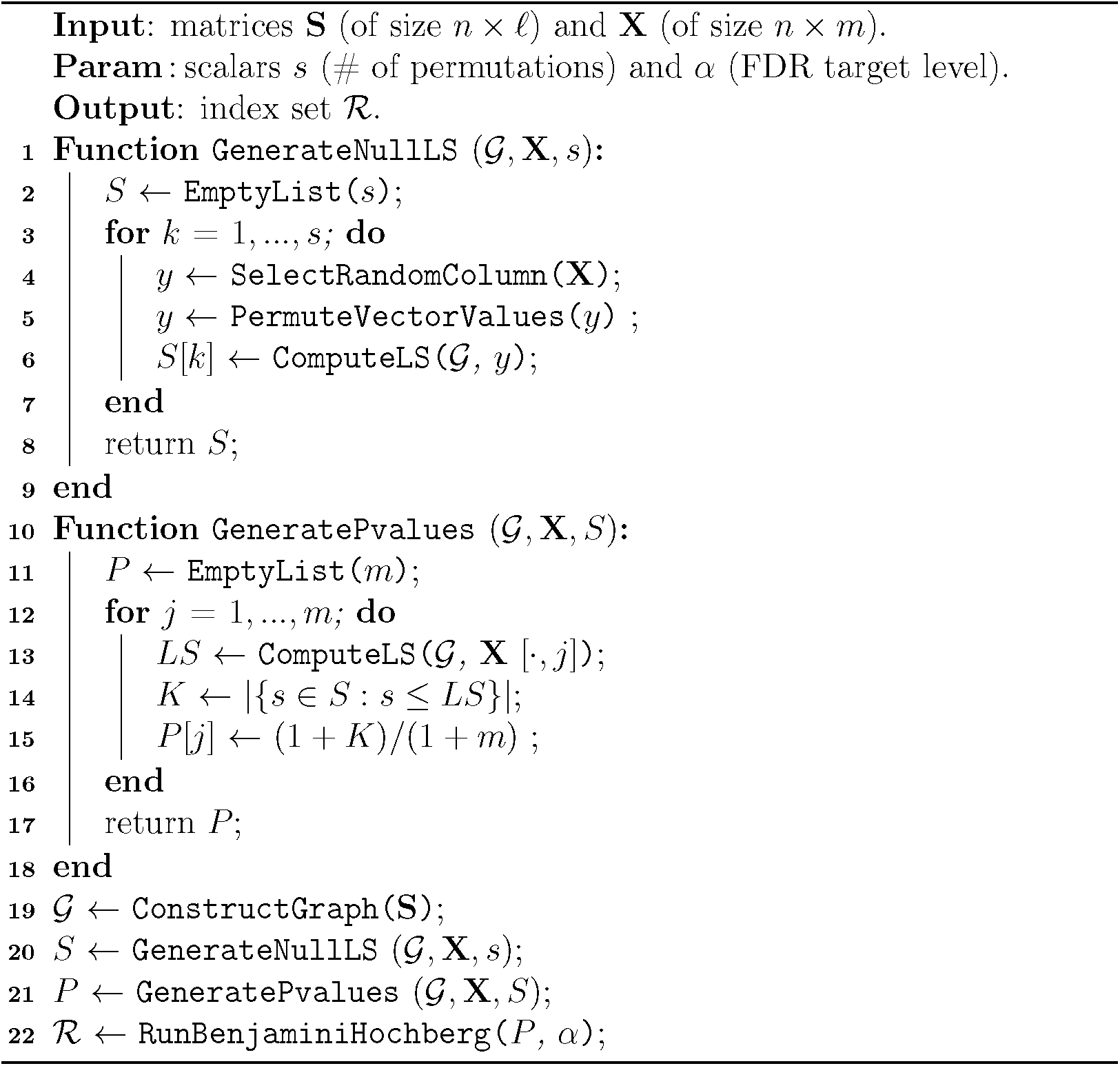

The inputs are two matrices: a “spatial” matrix **S** ∈ ℝ^*n×ℓ*^, and an expression matrix **X** ∈ ℝ^*n×m*^. During SV selection with the SGE data (Step Q in Figure 3), **S** and **X** correspond to the spatial coordinates and the gene expression data, respectively (Figure 2A); during SV selection with the scRNA-seq data (Step *(Q* in Figure 3), **S** ∈ ℝ^*N×t*^ and **X** ∈ ℝ^*n×m*^ correspond to the LM gene expression and the non-LM gene expression matrices, respectively (Figure 2B).

##### Computing the Laplacian Score

Let **x**_*j*_ ∈ ℝ^*n*^ correspond to the *j*-th column of **X** (i.e., the expression of gene *j*). We adopt the LS to measure the association between **x**_*j*_ and **S** for each *j* = 1, …,*m* as follows. We first construct a graph 𝒢 based on the pairwise distances among the *n* samples in **S**. By default, we construct a mutual nearest neighbor graph: each sample corresponds to a node, and there is an edge between two nodes if they are among the top *k* neighbors of each other in the ℓ-dimensional Euclidean space. Let **A** ∈ ℝ^*n×n*^ be a non-negative and symmetric adjacency matrix for G. The degree matrix is **D** = diag(*d*_1_,…, *d*_*n*_) with 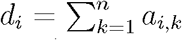, where *a*_*i,k*_ is the *(i, k*)-th entry of **A**, and *d*_*i*_ is the i-th row sum (or equivalently, the *i*-th column sum) of **A**. Then, the *graph Laplacian* is defined as **L** = **D** — **A**, and we use the (non-negative) LS to measure the relationship between **x**_*j*_ and **S** [27]:

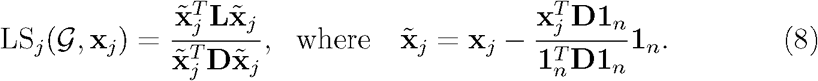

The variable 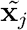 centers **x**_*j*_ and the denominator 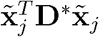 scales the score with respect to a discrete probability distribution on the *n* vertices proportional to their graph degree [27].

The LS is always non-negative and a low LS means a strong association between a gene expression and the “spatial” coordinates. This interpretation comes from its numerator in the quadratic form with the Laplacian **L**:

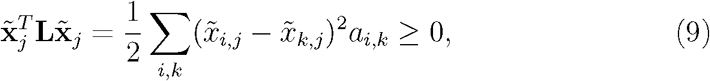

where *a*_*i,k*_, the non-negative entry from the graph adjacency matrix **A**, captures the spatial similarity between cell *i* and cell *k*, and the vector 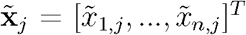 captures the expression of gene *j*. If two cells are spatially close, then *a*_*i,k*_ would be large and the difference in their expression 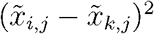 would contribute substantially to the score. If two cells are far apart, then *a*_*i,k*_ would be low and the difference in expression of gene *j* across them would contribute little to the score. An overall low LS indicates the gene expression in cells nearby are similar and the largest variation in expression occurs between cells that are far apart.

Thus, for a fixed graph 𝒢, a small LS indicates that there is an overall strong dependency between the gene expression values 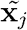 at each node and the similarity (or adjacency) matrix **A**.

*Remark:* In contrast to classical statistical measures, such as Pearson’s, Spearman’s, and Kendall’s correlations, which can only measure linear or monotonic relationships between univariate random variables, the LS offers two main advantages. First, the LS allows us to compare a univariate gene expression **x**_*j*_ to a multivariate spatial measurement **S**, by using them to construct a graph based on pairwise distances. Second, the LS can also capture non-monotonic or local associations based on comparisons between pairwise node value differences and their edge weights in Eq. (9). Moreover, we observed that its form in (9) is reminiscent of distance correlation [33], which has been a powerful means to capturing non-monotonic associations (Supp. Section A). From empirical simulations described later, we also find that LS is sensitive to monotonoic and non-monotonic local correlations, such as spatial expression hot spots in an SGE image.

##### Generating p-values from null Laplacian Scores

While LS was previously used as a scoring mechanism to prioritize graph-based features, we augmented it with significance testing for feature discrimination. We treat the LS of each gene as a test statistic and determine whether or not it is significantly small compared to observed LS values from genes that have no association with the graph structure. As such, we compute *p-values* for each gene based on its observed LS according to a permutation strategy with a fixed graph G constructed from **S** (Algorithm 1).

Given the *n* × *m* gene expression matrix **X** = [**x**_1_,…, **x**_*m*_], we first randomly sample a column/gene from **X** and then we permute its entries to obtain a vector **y**. For each of the *s* iterations, a *null* LS value is computed with a newly generated **y** and the same graph G (with **L** being fixed for all *s* iterations) based on Eq. (8), substituting **x**_*j*_ with **y**. The *s* null LS values represent the observed LS values when the gene expression is independent of the fixed spatial coordinates. For a given gene *j* and its score LS_*j*_ computed based on **x**_*j*_ in Eq. (8), we compute its p-value: *p*_*j*_ *= (K* + 1)/(*s* + 1), where *K* is the number of null LS values that are less than or equal to LS_*j*_. By default, we set *s* to be 10,000. When number of hypotheses is too large, however, we may obtain no rejections during multiple hypothesis adjustment, because the p-value threshold could be too stringent and fail to reject when *s* is not large enough. Therefore, if no rejections are observed due to too many hypotheses, we may also scale up the number of permutations until it is 100 times the number of hypotheses.

The permutation p-value counts the proportion of null LS values generated under the null hypotheses in Eq. (3) for SGE data and Eq. (4) for scRNA-seq data. The prescribed permutation procedure does not assume the data is generated from a specific distribution, nor does it rely on the asymptotic behavior of the test statistic. Thus, the procedure can simultaneously apply to the two data sources even if they have different noise distributions under Eq. (1) and Eq. (2).

*Remark:* Our pooling procedure leverages the facts that (1) the spatial coordinates and the Laplacian matrix **L** of each LS_*j*_ in Eq. (8) are identical, and (2) each LS_*j*_ is scale and shift invariant towards the expression of each gene **x**_*j*_. Instead of running our pooled procedure, one can compute the p-values based on constructing a null distribution for each gene by permuting itself *s* times, which, however, scales up the computational cost based on the number of genes. A sufficient condition for the pooling procedure to be valid is that the marginal distributions of each gene are identical after the normalization from **x**_*j*_ to 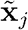in Eq. (8). We observed from simulated data that when realizations of **x**_*j*_ are randomly generated from different marginal distribution types (e.g., normal vs. uniform), their empirical null distributions of the LS are highly similar for each type when **L** is fixed. We also investigated on real data and found that the p-values from the pooled permutation approach are similar to that of computing permutations for individual genes (Supp. Section B.1).

##### Running the Benjamini-Hochberg Procedure

Given the p-values of each feature, one may consider choosing a fixed threshold to select non-null genes. However, the number of false discoveries could scale with the number of features tested even if all the features were null features.

Controlling the FDR for gene selection was put forward in early works for differential gene expression analysis in microarray and bulk RNA-seq studies [34, 35]. Like many of these works, we adopt the BH procedure [32] for its simplicity and theoretical guarantees of controlling the FDR. The BH procedure takes a set of p-values as input and selects a subset based on an adaptive threshold, such that FDR ≤ *α*, where *α* is the target level to control.

Although the BH procedure was originally shown to control the FDR under the assumption that the null p-values are independent [32], it has later been shown that the effect of dependence on FDR is negligible when the number of tests is sufficiently large for a general class of dependence [36, 35].

*Remark:* Recall that under the spatial model given in Eq. (2), the null hypothesis in Eq. (4) is true if the expression is not associated with the (unobserved) spatial coordinates. Because the LM gene expression 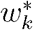 is a transformation of the spatial variable *s*_*k*_ and that its error vector 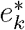 is almost independent of *s*_*k*_, a test for the association with 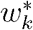 can be seen as more conservative than a test for the association with *s*_*k*_. Further, considering a non-SV gene (or null feature) with no error associated with the LM genes, its p-values can appear larger than the hypothetical p-value computed based on the association between *s*_*k*_ and *w*_*j*_. The same effect would apply to the p-values for the non-null features and make them appear more like null features, resulting in a loss of power to detect them.

Our procedure for selecting LM genes applies to different types of SGE data. One example is when only dozens or hundreds of genes are measured under FISH-based data, where our statistical test can determine how significantly the preselected genes vary spatially. Another example is when a tissue is spatially dissected into subregions and then RNA-seq is applied to each region. This approach can obtain tens of thousands of measured genes but at lower spatial resolution because only the average gene expression is measured within each subregion. Nevertheless, one can still apply our workflow by treating each region as a cell, and our multiplicity adjustment will guarantee control of the FDR for reproducibility, especially with a large number of genes to be tested. We do not require the cells or regions in SGE data to be further binned or aggregated. Instead, our graph-based approach directly captures the neighborhoods of the cells and can utilize the spatial information to the fullest extent. The removal of artificial binning within the SGE data further eliminates the need to bin the cells in the corresponding scRNA-seq data when we want to integrate the two data types.

#### 2.3.2 Latent Space Modeling

In order to reconstruct the spatial relationships among the cells in the scRNA-seq data, we rely on an updated spatial matrix, denoted as **Y** ∈ ℝ^*N×(t+r)*^, whose rows are the cells and whose columns include the LM genes and the selected SV genes. Then, we construct a graph and its Laplacian matrix, similar to how a graph and its Laplacian was constructed from the LM genes to compute the LS for SV gene selection (Section 2.3.1). We re-parameterize each cell based on spectral techniques similar to Laplacian Eigenmaps [37].

##### Spectral Construction

We reuse **L** ∈ ℝ^*n×n*^ to denote the graph Laplacian from graph 𝒢 constructed from **Y**. Let (*λ*_1_, **v**_1_), (*λ*_2_, **v**_2_), (*λ*_3_, **v**_3_),… represent the spectra of **L** in increasing order of the eigenvalues λ_*i*_. Each eigen-pair satisfies *λ*_*i*_**v**_*i*_ = **Lv**_*i*_ and can be successively solved based on the following optimization problems:

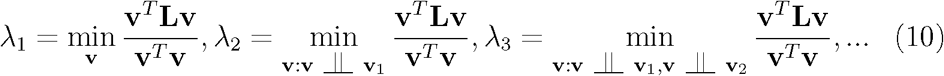

Here ╨ denotes linear independence, i.e., **x** ╨ **y** if and only if **x**^*T*^**y** = 0. To construct a *k*-dimensional embedding, we rely on the 2nd to the *(q* + 1)-th eigenvectors to generate an embedding matrix **Z** ∈ ℝ^*N×q*^, i.e., **Z** = [**v**_2_,…, **v**_*q*+1_]. Reusing the expansion in Eq. (9), we also have

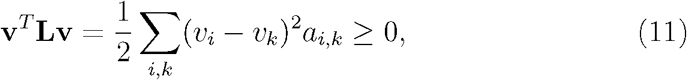

such that the minimizers of Eq. (10) embed two points at each embedding coordinate close to each other if their edge weight is large. The reason why we ignore the first eigenvector is due to the following known result.

###### Proposition 2.3. (The trivial eigen-solution)

Let 𝒢 be an undirected graph with non-negative weights. Then, for its Laplacian matrix **L**, we have the eigen-solution *λ*_1_ = 0 and **v**_1_ = **1** [26].

##### Trajectory inference

In the special case where *k* = 1, the solution to the problem is simply the second eigenvector, which can be seen as a trajectory that orders the cells (Figure 3). Formally, solving for the second eigenvector is equivalent to solving:

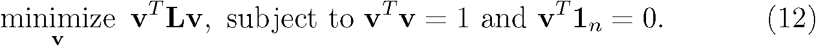

The reformulation comes from first turning the denominator of the objective into a constraint **v**^***T***^**v** = 1 in Eq. (10), and then applying the equivalence **v** ╨ **v**_1_ ⟺ **v**^*T*^ **1**_*n*_ = 0 for the second problem in Eq. (10).

Interestingly, the two constraints in Eq. (12) ensure that the solution is scale and shift invariant. We leverage this observation to derive a theoretical result regarding the uniqueness of the cell orderings under the continuity assumptions in Section 2.2.3.

###### Theorem 1.

*Let 𝒢 be a connected graph constructed from data points sampled from a continuous distribution, then the solution to Eq*. (12) *generates a unique ordering of the data points*.

*Proof*. The proof is given in Supp. Section C. □

*Remark:* When the data is generated from a continuous spectrum, well-established clustering techniques cannot exploit inter-cluster and intra-cluster relationships between the points. We argue that constructing orderings of cells is itself a powerful approach to interpret continuous expression gradients along fine-grained structures. A variety of TI methods have been used on scRNA-seq data to extract pseudo-temporal ordering of cells [25]. However, these methods rely on all the genes to construct their latent variable, whereas `GLISS` only relies on the SV genes. The differences in the inferred trajectories can be substantial when the signal is limited to a few spatial genes. While existing TI methods can substitute `GLISS`’ routine after selecting SV genes, we selected the current graph-based embedding framework because of its consistency with the LS approach used for feature selection (Section 2.3.1), as well as its uniqueness guarantees based on Theorem 1. Additionally, `GLISS` can extend to multi-dimensional latent variables by solving additional eigenvectors, whereas most TI methods only provide 1-D (temporal) orderings.

#### 2.3.3 Gene Dimension Reduction and Clustering

`GLISS` fits a spline model as a function of the latent structure to obtain lower-dimensional characterization of each gene. Then it utilizes the lower dimensional space to cluster the genes into groups that share similar spatial patterns.

##### Dimension reduction

Following the latent space inference setup (Section 2.3.2), we use the embedding matrix **Z** ∈ ℝ^*N×q*^, based on the solutions of Eq. (10), and the expression matrix **Y** ∈ ℝ^*N×(t+r)*^ as inputs^4^, and fit a spline model with *s* degrees of freedom, such that each gene is represented as the columns within a gene embedding matrix **H** ∈ ℝ^*s×(t+r)*^. For each gene, we fit a natural cubic spline model given by

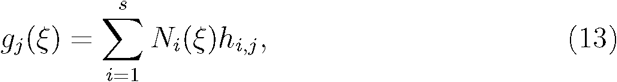

where *N*_*i*_(*ξ*) is the i-th basis function from the family of splines. *h*_*i,j*_ represents the *(i, j*)-th entry of **H** obtained from solving a generalized additive model [38]. We can then visualize the de-noised continuous expression along the inferred spatial structures.

*Remark:* Our dimension reduction technique is specific to the inferred latent structure because it uses the parameterization from the fitted models with the latent structure as the predictor variable. It also provides a strategy to handle the case when genes are correlated due to the noise but not due to the spatial variable.

##### Reducing noise correlation across genes

We simulated data generated along a 1-D latent variable (Figure 4): we ordered the cells in 1-D, and then generated the expression of SV genes under the model under Eq. (7) with six different spatial patterns (including monotonic and non-monotonic ones). The six groups of SV genes with distinct spatial patterns are plotted in order as columns in the heat map; the remaining non-SV genes are generated via Gaussian noise with five correlated groups, whose columns are also plotted in order (Figure 4A). Note that the non-SV genes do not appear to be associated with the cell ordering. When computing the gene-to-gene similarly matrix in the *N*-dimensional space (Figure 4A), one would observe groups of genes that are highly similar including the non-SV gene groups that are similar due to the noise correlation. After fitting the splines for each gene from the original data, the fits that correspond to the same spatial pattern will share similar spline coefficients, whereas the coefficients for these non-SV gene groups will have coefficients that are all close to zero (Figure 4B). The influence of the correlated noise can be mitigated as the inferred 1-D spatial variable was used to reduce the dimension of each gene when computing the gene-to-gene similarity (Figure 4C): all the null features across the five groups appear to look more similar to each other with respect to the latent structure, whereas the six groups of non-null features each appear to be more distinguishable from the similarity matrix.

**Figure 4:**
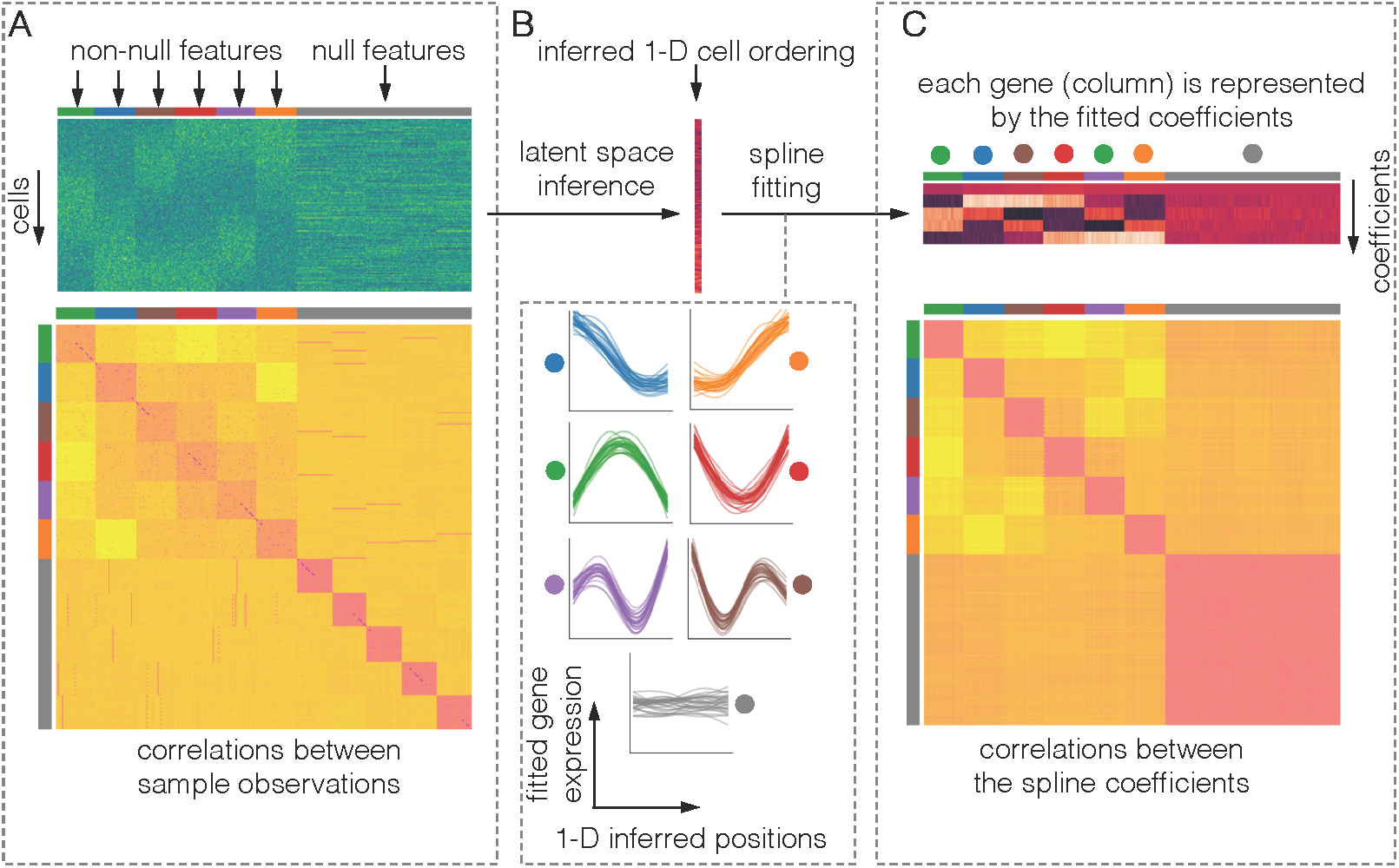
Two approaches to learn gene-to-gene similarities. (A) Gene-to-gene similarities are computed from the the original gene expression. (B) The cells are embedded into a latent space, and each gene is fit against the latent variable. (C) Gene-to-gene similarities are computed based on each gene’s fitted coefficients. Using the coefficients reduces correlation unrelated to the spatial variation.

##### Gene clustering

We can then utilize the new gene embeddings to identify groups of genes that share similar spatial patterns. Not to be confused with the spectral approaches in inferring latent *cell* positions, we apply spectral clustering [26], treating each *gene* as a sample represented by its vector of coefficients in the matrix **H** solved from Eq. (13). While spectral clustering is the default for `GLISS`, it also would work with other clustering approaches as long as they take **H** as an input. Because the spatial information is already captured by the coefficients and the non-spatial effects are already mitigated in **H**, we expect the clustering quality for these approaches to improve over naively clustering the genes from the raw expression matrix (Figure 4).

*Remark:* As the number of SV genes reach the order of hundreds or thousands, grouping genes with similar spatial patterns allows one to investigate the enrichment of specific pathways or processes. We will show from simulated and real data analyses that `GLISS` not only eliminates the need to discretized the cells into bins and average over them, but it can even leverage the continuous spatial information to improve clustering and interpretation.

## 3 Results

In this section, we describe the simulation and real data analyses that assess `GLISS`:

1. We first evaluate the initial step of selecting LM genes using simulated SGE data (Section 3.1). We benchmark our graph-based feature selection procedure based on the LS (Section 2.3.1) against existing SV detection methods (`SpatialDE` [20] and `scGCO` [22]) in terms of power and FDR (Section 2.2.2).
2. Then we re-analyze two real SGE datasets. Because we do not have the ground-truth SV genes to compute power or FDR, we focus on how the selected SV genes overlap across different methods or technical replicates as a measure of replicablity.
3. We then simulate scRNA-seq data with LM genes and correlated noise across all genes to mimic mixed spatial and non-spatial effects in the data. In addition to evaluating the SV detection quality, we assess the the subsequent steps of latent space reconstruction (Section 2.3.2) and gene dimension reduction (Section 2.3.3). In particular, we compare our inferred latent structure with that constructed from existing TI methods (`Palantir` [39] and `PAGA` [40]) that do not utilize the LM genes like `GLISS`.
4. Finally, we evaluate `GLISS` on real scenarios by re-analyzing the data from two high-profile integrative analysis studies for the liver [3] and the intestine [4], respectively (Section 3.4). We aim to demonstrate that `GLISS` can produce results consistent with the original studies, but with a much simpler and more reproducible workflow. In particular, the two original integrative studies relied on inferring the discretized spatial zones of the cells in their customized pipelines to determine new SV genes. Thus, we not only want to demonstrate that our novel SV genes highly overlap with theirs, but also show that our continuously reconstructed latent ordering is similar to their binned ordering. Additionally, we identified SV gene sets from the two studies using `GLISS`’ gene dimension reduction and clustering workflow and validated newly discovered gene sets with the Gene Ontology (GO).

### 3.1 SV detection in Simulated SGE Data

To begin, we simulated SGE data alone with both spatial coordinates and gene expression (Figure 2A). We generated true null and non-null features according to the model in Eq. (1), with different spatial patterns for the non-null features. We then computed the statistical power and FDR (Section 2.2.2) of `GLISS`’ graph-based feature selection procedure, benchmarked against those of `SpatialDE` and `scGCO`, for each spatial pattern.

For the simulation, we re-used the 2-D spatial coordinates from a MER-FISH dataset with 1,045 cells [41]. For the gene expression matrix, we used 200 genes to mimic the number of genes measured in the SGE data. Each simulation used a given expression pattern with 4 different resolutions (from global to local, Figure 5A) to generate 100 SV genes (i.e., non-null features), with 25 genes per resolution. The remaining 100 genes are non-SV genes (i.e., null features). Based on Eq. (1), we defined *fj(si)* to be 0 for all *i* if gene *j* is a null feature. For non-null features, we assigned non-constant functions fj *(si)* in 4 different simulations, each associated with a particular spatial expression pattern (Figure 5A):

- Linear: the SGE monotonically increases diagonally;
- Quadratic: the SGE quadratically decreases from a diagonal;
- Radial: the SGE decreases gradually from the origin;
- Sinusoid: the SGE is a 2-D sinusoid function a specific frequency.

**Figure 5:**
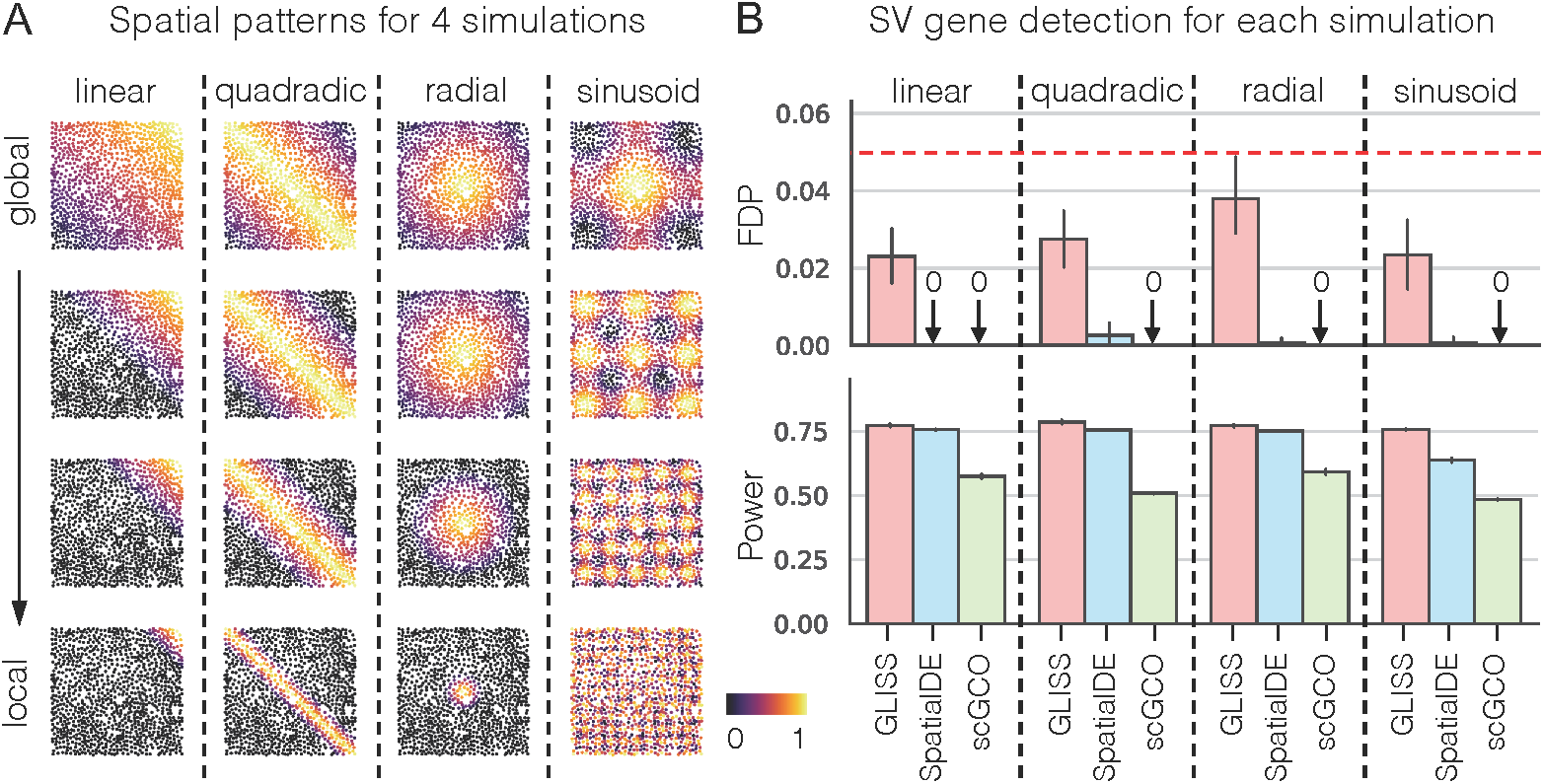
SGE simulation. (A) Different types of spatial gene expression patterns within cells organized in a 2-D coordinate system. Each simulation consists of 100 non-null features, with 25 from each resolution. (B) FDP and Power comparison across `GLISS`, SpaialDE, and `scGCO`. Each group corresponds to a simulation with a specific spatial pattern repeated over 20 trials. The target FDR level is set to be 0.05 for these methods.

We repeated 20 trials per simulation, each with independent Gaussian noise to assess sensitivity of the SV gene detection methods.

We observed that `GLISS` is at least as powerful as `SpatialDE`, the state-of-the-art method for SGE analysis (Figure 5B), and notably more powerful than `scGCO`. Breaking down the power of each simulation into the 4 resolutions (Extended Figure 12), we confirmed that all methods had the most difficulty selecting genes with highest resolution of each spatial pattern. However, `GLISS` was able to occasionally select a few high resolution genes when other methods did not detect any (Extended Figure 12, last row). Most noticeably, `GLISS` outperforms on the sinusoid example, where the signal has high frequencies and there is also high local variability in the data. The gain in statistical power for `GLISS` was accompanied by higher FDP compared to the other methods, but the level was controlled under the target level of 0.05 on average (Figure 5D).

### 3.2 SV detection in Real SGE Data

Next, we considered two real SGE datasets that were originally analyzed by `SpatialDE` and `scGCO`. Because no ground-truth SV genes were available for these datasets, we mainly investigated the gene set overlap between `GLISS` and other methods:

- The mouse olfactory bulb (MOB) dataset [15], including 3 replicates sampled from separate regions of the same tissue.
- The breast cancer (BC) dataset [15], which had previously been analyzed by `SpatialDE`, `scGCO`, as well as `trendsceek` [21].

The 3 replicates in the MOB data had been previously analyzed by `SpatialDE` and `scGCO`. We first re-evaluated the gene set overlaps across each method for each replicate, and included the overlapping gene sets identified by `GLISS` (Extended Figure 13A). We noticed that the number of SV genes selected by `GLISS` was similar to that of `SpatialDE`, and much less than that of `scGCO`, which indicated that `GLISS` was not over-selecting. Next, we assessed how the SV genes were replicated for each method. The overlap score we defined is the intersection of all gene sets divided by the union of them (which generalizes the Jaccard similarity score). While `GLISS` selected fewer genes compared to other methods, the gene overlap among the technical replicates was higher in `GLISS` (Figure 6A).

**Figure 6:**
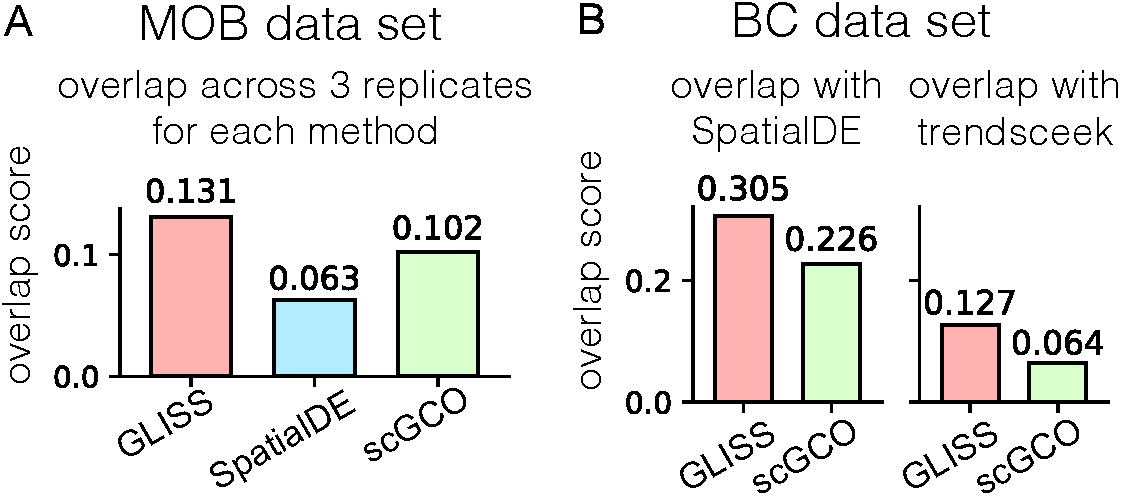
(A) The mouse olfactory bulb (MOB) SGE dataset [15]. Each method is evaluated based on the overlap across 3 gene sets, each corresponding to the selected SV genes from a replicate. (B) The breast cancer (BC) SGE dataset [15]. We use the gene sets from `SpatialDE` and `trendsceek` as two references, and compare how the gene sets of `GLISS` and scGSCO overlap with each reference respectively.

On the BC dataset, we used the SV genes from `SpatialDE` and `scGCO` as two independent reference gene sets, and evaluated how the newer methods, `scGCO` and `GLISS`, overlapped with each reference. The two references, `trendsceek` and `SpatialDE`, were very different from each other because the former selected 15 genes whereas the latter selected 115 genes, with 10 overlapping genes (Extended Figure 13B). When comparing `scGCO` and `GLISS` against these two different references (Extended Figure 13B), we found that `GLISS` had better overlap with both `trendsceek` and `SpatialDE`, even though `GLISS` selected fewer SV genes compared to `scGCO` (Figure 6B).

### 3.3 Performance on Simulated scRNA-seq Data

Having demonstrated `GLISS`’ ability to detect SV genes in SGE alone, we next assessed how well it selects new SV genes in scRNA-seq data during integrative analysis. To mimic the liver study [3] (Section 1), we simulated data consisting of a true latent variable, representing the position along the central-portal axis (Figure 1A), expression of LM genes and the non-LM genes measured in scRNA-seq (Figure 2B). Given the previous SGE simulation, we assumed that some LM genes were obtained from SGE data to focus on the outcome of the integrative workflow. The non-LM gene expression could be further divided in into a non-null feature matrix corresponding to the SV genes, and a null feature matrix (Figure 2B).

Mimicking the previously mentioned hepatocyte study [3], we generated 1,500 cells and 6,000 genes. There are a total of 906 SV genes, 6 of which we assume to be known as LM genes. The LM genes consist of 3 templates, with genes per template, and the SV genes consist of 6 templates, with 150 genes per template (Figure 7A). The first three SV gene templates resemble the 3 LM gene templates (Supp. Section D). The remaining 5,094 null features (non-SV genes) were generated to be independent of the latent coordinates *ξ*_1_, …,*ξ*_*N*_. To generate noise correlations across the genes (including LM genes, SV genes, and non-SV genes), we generated the noise vectors **e**_1_,…, **e**_*N*_ (with **e**_j_ = [*e*_1,*j*_,…, *e*_*M,j*_]^*T*^), such that each vector was independently sampled from a multivariate normal distribution with diagonal blocks in (the gene-to-gene) covariance matrix: each block was composed of 150 features with within-block pairwise correlation equal to *ρ* (Extended Figure 14). Additional details are provided in Supp. Section D.

**Figure 7:**
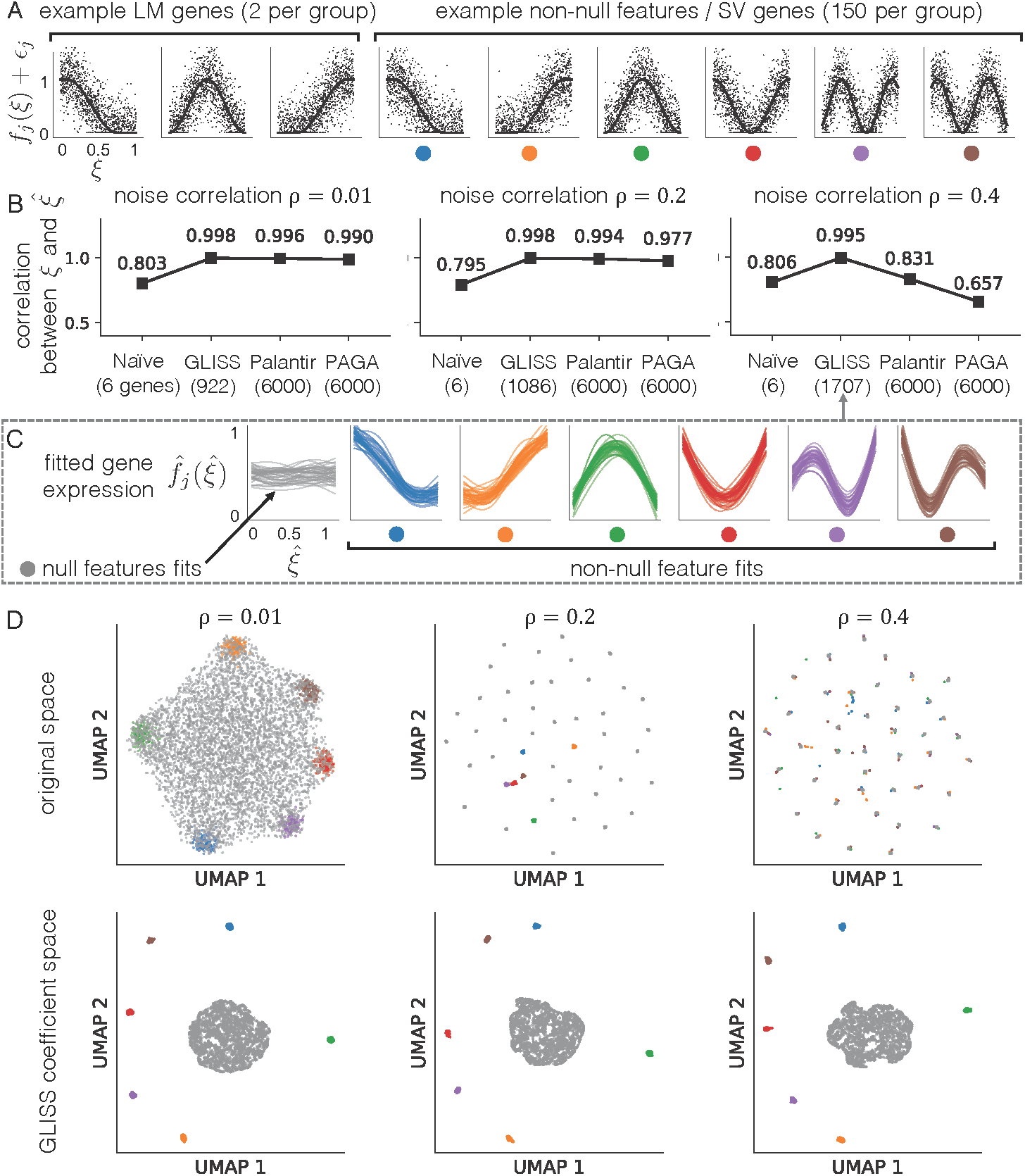
Simulation results for scRNA-seq data. (A) Examples of the of the LM and the non-LM SV gene expression, as a function of the ground-truth latent cell ordering. The 6 non-null templates are indicated by the six colored markers. (B) The absolute Spearman’s correlation between the true *ξ* and the inferred variable 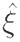. The subplots correspond to simulations with different noise correlation. (C) The spline fits with *ρ =* 0.4 grouped by ground-truth non-null groups in (A). (D) The UMAP embedding of all the genes. Each grey dot corresponds a null feature, and the other dots are non-null features colored by their template.

Given the ground truth latent variable *ξ* (e.g., the radial position), we assessed the latent variable recovery based on the absolute Spearman’s correlation between *ξ* and its estimate 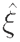. We evaluated two ways to obtain 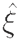: (1) the 1-D graph embedding using both the LM genes and the inferred SV genes from the pipeline, according to Section 2.3.2 (`GLISS`) and (2) the 1-D graph embedding only using the LM genes (Naive). We benchmarked `GLISS` against `Palantir` [39] and `PAGA` [40], two state-of-the-art unsupervised approaches that use all genes to infer 1-D cell trajectories. Even though all methods obtain near perfect latent space reconstruction under little correlation, i.e., *ρ =* 0.01, `GLISS` outperformed both `Palantir` and `PAGA` noticeably under higher noise correlation, i.e., *ρ* = 0.4, (Figure 7B). The Naive approach recovered *ξ* with lower correlation because it only uses the LM genes and the noise in the LM genes obscures the true cell ordering to some extent. Not only did `GLISS` reconstruct the trajectories accurately despite noise correlations, it also selected the SV genes with high power and controlled FDP^5^ as seen in the SGE simulations (Extended Figure 15A). On the other hand, we found that `Palantir` and `PAGA` are more sensitive to the correlation structure in the noise when they use all available genes as `GLISS` does. Both `Palantir` and `PAGA` improved their estimates to the same level as `GLISS` when their input matrix only included the LM genes and SV genes selected by `GLISS` (Extended Figure 15B). This result indicates that a major advantage of `GLISS` is its ability to select SV genes by leveraging information from the LM genes, which other current methods do not do.

Furthermore, we evaluated `GLISS`’ gene dimension reduction approach (Section 2.3.3) with the same simulation setup and compared it to directly computing the gene-to-gene similarities in the original space without SV gene selection or dimension reduction (Naive). `GLISS` was able to fit distinct spline fits to each of the six non-null feature groups that resembled the pattern we generated, such that the coefficients of the within-group features would be more similar to that of across-group features (Figure 7C). We applied UMAP [42] to visualize all genes (including SV and non-SV genes) in two different ways: (1) using the coefficient matrix output from `GLISS`, and (2) using the original expression matrix for the Naive approach (Figure 7D). When the noise correlation is large (e.g., ρ = 0.4), the naive approach aggregated gene groups based on their noise correlation, resulting in distinct clusters of null features (colored gray) with the non-null features (shown in six other colors) hidden among them. Meanwhile, the correlation had almost no noticeable influence on `GLISS`, which computes gene-to-gene similarities based on the smoothing coefficients instead of the original noisy expression. We repeated the UMAP visualization of only the SV genes (selected by `GLISS`) and found that the segregation of different non-null templates was still more obvious for `GLISS` (Extended Figure 15C).

In this simulation, however, the noise among LM genes are correlated with that of some null features, which violates the condition of Proposition 2.2. `GLISS`’ selected these features under strong noise correlation of *ρ =* 0.4, because each of them had a small enough p-value corresponding to their null hypothesis in Eq. (4). When such genes are selected in real scRNA-seq data, we may not be able to distinguish them from SV genes. However, `GLISS`’ gene dimension reduction could potentially isolate this group of null features based on their spline coefficients: we observed that the null features are grouped away from the non-null features despite being selected as SV genes in the UMAP space.

### 3.4 Integrative Analysis on Real Data

The last part of our results is the real data analysis of the integrative workflow with two systems: hepatocytes in the liver (previously studied by [3]) and enterocytes in the intestine (previously studied by [4]). The goal of our re-analyses is to re-identify the SV genes discovered in the original pipeline with our simplified workflow and determine if we can discover new SV gene sets that can be validated via the Gene Ontology (GO).

#### 3.4.1 Hepatocytes in the Liver

We analyzed the hepatocyte dataset from study [3], which included data from both smFISH and scRNA-seq. We focused on two aspects in this analysis: (1) comparison of different approaches that infer latent cell positions, and (2) investigation of the resulting expression patterns of specific LM and SV genes identified from the original study. The original study selected six LM genes from smFISH and then developed a Bayesian model on the six genes to specifically infer the spatial locations (based on nine discrete zones) of the cells from scRNA-seq. We used the same LM genes when analyzing the scRNA-seq data (with 1,415 cells and 8,883 genes, Supp. Section E.1), which also made the scRNA-seq analysis comparable for this example.

There are three types of inferred 1-D latent positions we compared (Figure 8A): (i) Original and Naive estimates that only use the 6 LM gene expression; (ii) `GLISS` which uses the LM genes and the selected SV genes, with a total of 1483 genes; and (iii) `Palantir` and `PAGA` which are unsupervised and consider all 8,883 genes (Figure 8A). The correlation between the original and the Naive was relatively high, even though the original model in [3] required extra information of the gene expression from the smFISH data to map the cell positions. `GLISS` was more correlated with the Naive approach because the SV genes were chosen based on their association with the LM genes. `Palantir` and `PAGA`’s estimates were more correlated with each other and less correlated with the other estimates, which indicates that non-spatial signals may exist in some of the genes that were not selected by `GLISS`.

**Figure 8:**
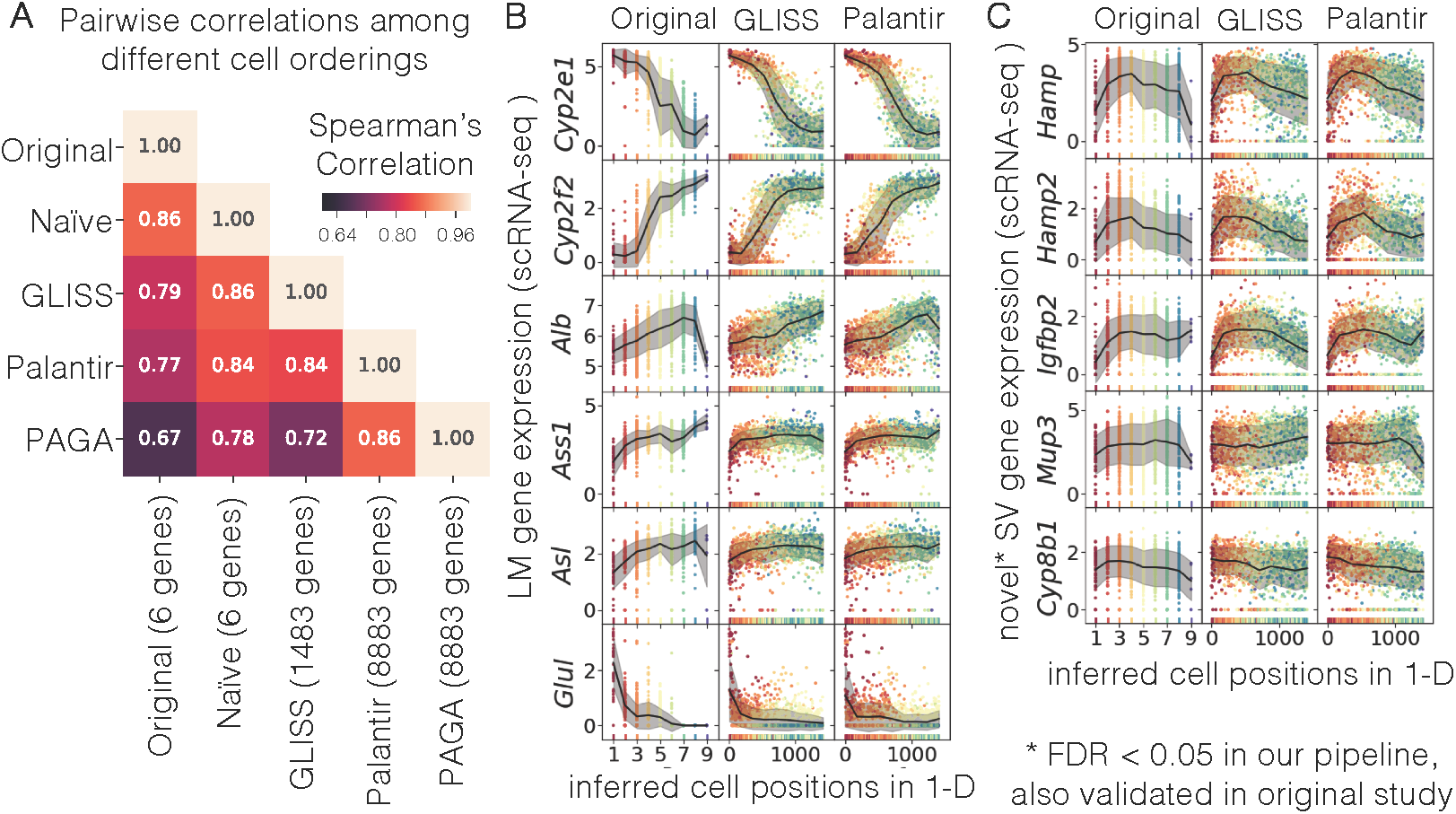
Real data analysis for hepatocyte data. (A) Pairwise correlations of the 1-D latent variable inferred by different methods. The number of genes in the parentheses indicate the ones used for latent space inference. (B) LM genes from scRNA-seq data. (C) Previously validated SV genes from the original study. (B-C) Each row in the plotted grid corresponds to a different gene and each column corresponds to a different method that infers the 1-D cell ordering which was compared in (A). Each gene expression is plotted against the inferred cell position in 1-D by different methods.

Given the different inferred cell positions, we investigated the spatial patterns of specific genes as a function of the positions. First, we plotted the expression of the LM genes (Figure 8B). The plots of the same LM genes (each row) highlight some discrepancies among the latent variables. For instance, the positions of cells with discretized positions (Original) have less resolution than the those with continuous positions (`GLISS` and `Palantir`) along 1-D spectrum. Interestingly, the expected monotonic behavior of *Cyp2e1, Ass1*, and *Asl* (based on the smFISH data in the original study) appeared to be non-monotonic by Original and `Palantir`, compared to `GLISS` (Figure 8B). Next, we considered 5 novel non-monotonic genes (Figure 8C): *Cyp8b1, Mup3, Igfbp2, Hamp2*, and *Hamp*, which were identified and validated in the original study. To declare a SV gene as non-monotonic, the original analysis required the average zone expression to peak between zones 3 and 7, and that this peak value had to be 10% higher than the maximal value between zones 1 and 9. `GLISS` was also able to identify these validated SV genes, but with its much simpler routine. In fact, `GLISS` identified additional genes that were similarly non-monotonic using our gene dimension reduction and clustering workflow. We found that a cluster (with 46 genes) which included *Hamp* and *Hamp2* was enriched in “alpha-amino acid metabolic process” (FDR < 0.1). The original study mentioned that they were not able to identify GO terms enriched with their non-monotonic genes [3], but we hypothesize that their definition of non-monotonic genes might have been too specific and led them to filter out other genes that could have been identified otherwise.

#### 3.4.2 Enterocytes in the Intestine

We next analyzed enterocyte data to investigate spatial expression patterns along the villus-crypt axis (Figure 1B). In a recent study [4], the authors characterized landmark genes with bulk RNA-sequencing of laser capture microdissected tissue (LCM-RNA-seq). Then, they linked their LCM-RNA-seq SGE data with scRNA-seq data from in a different study [43] to identify novel SV genes. The focus of this dataset is to compare the intermediate results from each component of `GLISS` with theirs.

After running our workflow, we found good agreement between the selected gene sets between `GLISS` and the original study (Figure 9A). Given the SGE data, which consisted of bulk RNA-seq of 6 consecutive regions, `GLISS` selected over 350 out of 1,005 as LM genes. Then, it selected 7,274 out of 9,656 genes from the scRNA-seq data (with 1,383 cells), among which 1,920 high variance genes were used for gene dimension reduction and clustering (Supp. Section E.2). Although `GLISS`’ LM genes overlapped with those from the original study by 63, the SV genes detected from the scRNA-seq data overlapped significantly more.

**Figure 9:**
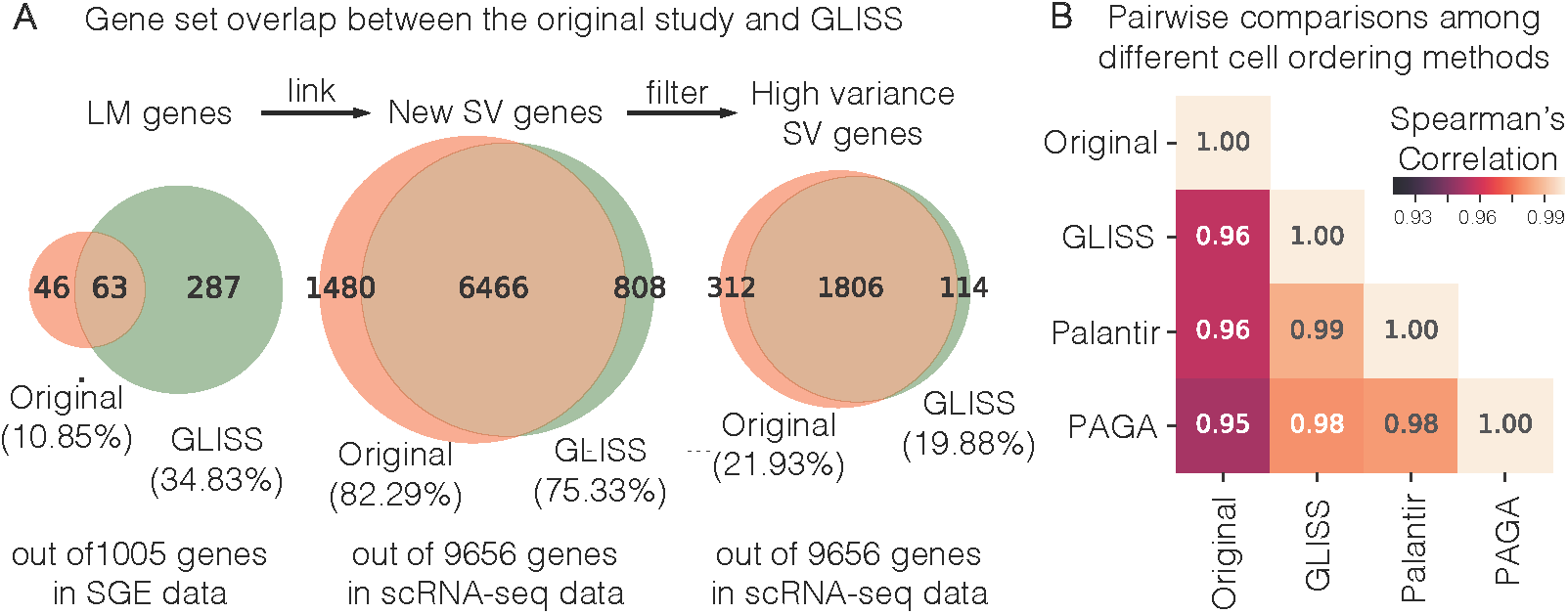
Real data analysis for hepatocyte data. (A) The venn diagrams comparing the overlap between the LM genes, SV genes, and filtered SV genes from `GLISS` and the original study. (B) Pairwise comparisons of the inferred 1-D latent variable, similar to that of Figure 8.

By assessing the gene sets, we reasoned that as long as the overlapping LM genes contain similar spatial information, two methods can still end up selecting highly overlapping SV genes in scRNA-seq data. In fact, we also observed high correlation among the latent cell positions inferred by different methods (Figure 9B). The number of genes used by the original study was on the order of hundreds; the number novel SV genes from scRNA-seq data used by `GLISS` was on the order of thousands; and all genes were used by `Palantir` and `PAGA`. Thus, it is not surprising that TI methods (`Palantir` and `PAGA` which used all genes) had high concordance with the original and `GLISS`’ inferred cell orderings. Indeed, the most common use case for TI methods is to identify differentiation signatures when the majority of genes vary in the same way.

Last, we compared the gene clusters identified in original studies with the clusters identified by `GLISS`. The original study represented each gene with 7 features which led to 5 clusters, whereas `GLISS` represented each gene based on 13 features from the fit coefficients (10 knots, natural cubic splines) and identified more clusters (Supp. Section E.2). We visualized both high-dimensional representations by projecting each gene into a 2-D space (Figure 10). `GLISS` exhibited complex structures corresponding to gene clusters that were difficult to identify in the the original 7-dimensional data: Cluster 3 identified by `GLISS` (red, 60 genes) was enriched in “ATP metabolic process” and other related processes (FDR < 0.1), but the genes from Cluster 3 appeared to be hidden in the original gene embeddings. Cluster 6 identified by `GLISS` (pink, 33 genes) was enriched in “cellular organization” and other related processes (FDR < 0.1). Even though the genes in Cluster 6 were visible in the original gene embeddings, this cluster would have been too small to be detected in the original study.

**Figure 10:**
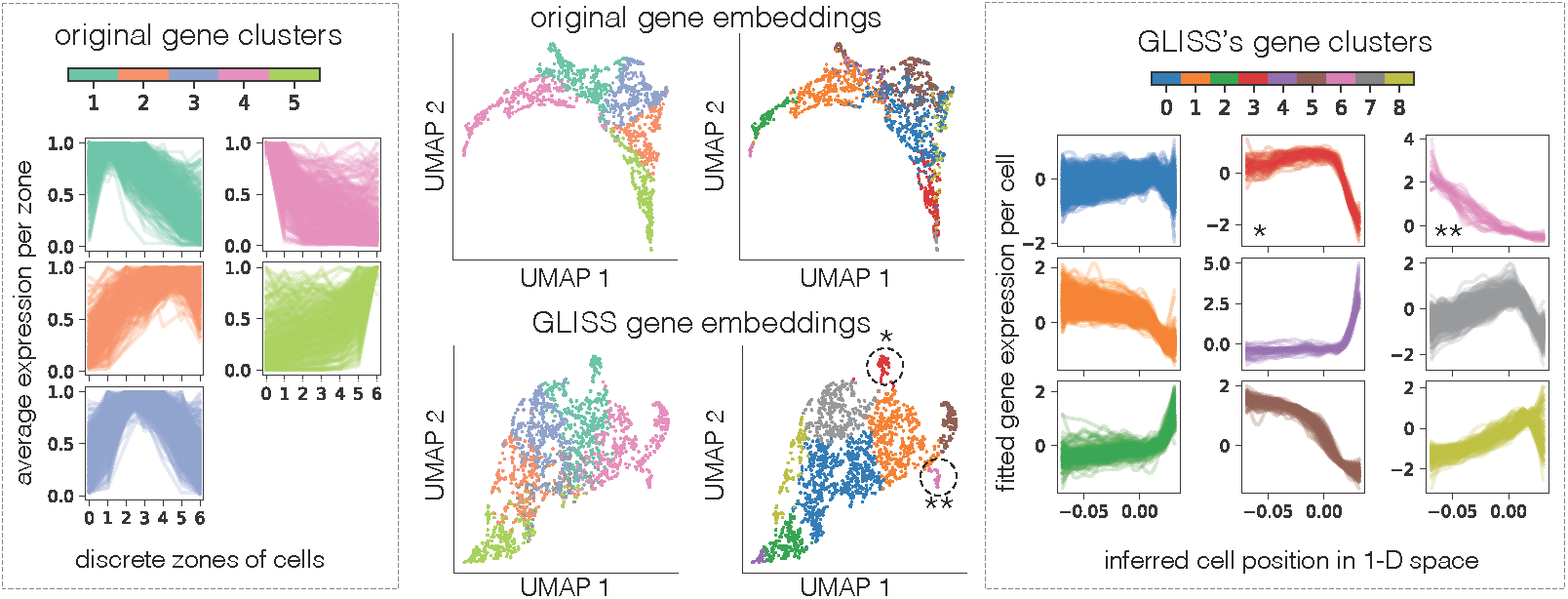
Gene clusters from the enterocyte study. Colors of the lighter shade (used in the left panels) correspond to the clusters from the original study; colors of the darker shade (used in the right panels) corresponds to the clusters identified by `GLISS`. The middle plots are UMAP embeddings of the SV genes using the original gene expression matrix (top row) vs. using our gene embeddings from the coefficient matrix (bottom row). The points in each column of the middle plots are colored by the cluster IDs shown in the legend of the left and right dotted boxes. The left and the right panels in the dotted boxes plot the groups of averaged (left) or fitted (right) gene expression profiles against the spatial variables by the original study (left) and `GLISS` (right). Cluster 3 and Cluster 6 are marked in the middle plot and the right box with ∗ and ∗∗.

## 4 Discussion

GLISS is a novel framework that integrates spatial gene expression data with scRNA-seq data to simultaneously select spatial gene features and identify hidden spatial cellular structures. It relies on a graph-based feature selection method that is sensitive to non-monotonic associations to determine spatial genes, so that one can obtain a rich repertoire of gene expression patterns for downstream analysis such as gene clustering. As more SGE data and scRNA-seq become available, there will be growing interest in performing integrative data analysis, especially to understand cell types and states with spatial information. Applying `GLISS` to these new datasets would require little parameter adjustment thanks to its simplified workflow. Our statistical components were built with reproducibly and computational efficiency in mind, so researchers can follow up with the newly discovered SV genes in future studies. `GLISS` can be combined with existing pipelines and our data preprocessing procedures were handled by using a standardized library for scRNA-seq data called scanpy [29]. On the other hand, we still had to rely on customized data quality preprocessing pipelines specific to each study for the SGE data because not all of them have been standardized. In the future, it would be important to systematically investigate how sensitive the integrative analysis is to the the SGE data preprocessing so we can build more confidence about the LM genes.

## 5 Acknowledgments

We would like to thank Trevor Hastie for helpful feedback and discussions.

## A The Laplacian Score as an Association Measure for Non-monotonic Relationships

To detect non-monotone relationships, many methods, including the distance correlation [33], have been proposed to directly test the hypothesis of independence *H*_*0*_ : *F*_*X,Y*_ = *F*_*X*_*F*_*Y*_, i.e., the joint distribution of two random variables is the product of their marginal distribution. Given paired samples 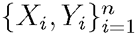 from two random vectors, the original distance dependence statistics (i.e., distance covariance) takes on the form of

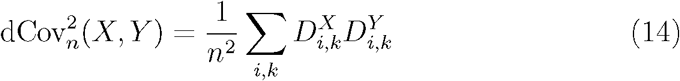

with

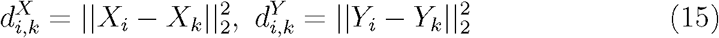

and

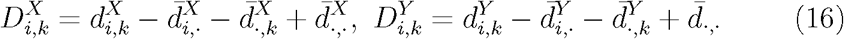

where 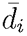,. is the i-th row mean, 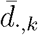 is the *k*-th column mean, and 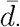,. is the grand mean of the pairwise distance matrix whose (*i,k*)-th is *d*_*i,k*_. In other words, the distance correlation first considers *n × n* Euclidean distances between each pair of data points for each variable, and then only compares the two n x n distances matrices distance matrices instead of the n original data pairs. If one vectorizes the two distance matrices, the distance covariance would be the inner product of the two vectorized matrices (after row, column and grand-mean centering). It can be shown that the distance covariance is always greater than or equal to zero, and almost equal to the population value of the distance covariance with a large number of observations; further, the population value is equal to zero if and only if the two variables are independent [33]. For an index pair *(i,k)*, if ‖*X*_*i*_ — *X*_*k*_‖^2^ is relatively large compared to other index pairs, then increasing ‖*Y*_i_ — *Y*_*k*_‖^2^ would dramatically increase the overall distance covariance. Essentially, the distance covariance measures the concordance between the pairwise dissimulates.

The LS also compares the pairwise relationships between variable *S* and variable *X*_*j*_, where *S* is multivariate and *X*_*j*_ is univariate. Based on the reformulation in Eq. 9, the term (*x*_*i*_ — *x*_*k*_)^2^ measures the dissimilarity between the points in the 1-D Euclidean space, whereas *a*_*i,k*_ measures the spatial similarity instead of dissimilarity. Thus, one main difference between the distance covariance and LS is that they differ in their direction when interpreting the strength of associations. Another way to interpret the LS is based on a specific graph where each edge follows 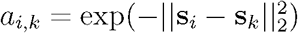. The continuous and bijective map between 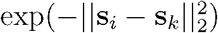 and 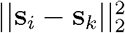 indicates the equivalence between the products *a*_*i,k*_(*x*_*i*_ − *x*_*k*_)^2^ (LS) and 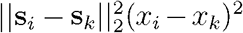 (distance covariance) in terms of their contribution towards the their respective association measure. Other measures such as certain kernel-based measures have been shown to be equivalent to distance covariance due to similar product forms and bijective transformations [44, 45, 46]. Similarly, the LS and distance covariance can achieve similar sensitivity towards non-monotonic relationships with an appropriate kernel transformation of the distance covariance or modifications of the graph edge weights in the LS. In contrast to the aforementioned methods, classical correlation measures are not able to determine such non-monotonic relationship due to their construction of directly comparing the paired observations instead of the pairwise distances. For instance, Pearson correlation only computes the inner product of the two univariate random variables, but negative terms can cancel positive ones when computing the inner product (e.g., due to symmetry) and can make the Pearson correlation appear to be close to zero when the two variables are not independent.

## B Implementation Details

### B.1 Pooled Permutation

While implementing the Algorithm 1, we implemented graph Laplacian Score (LS) and the Benjamini-Hochberg (BH) procedure ourselves due to their simplicity. We set the FDR target level to be *α =* 0.05 for the simulations as well as the real data analysis. By default, we set the number of permutations *s =* 10,000 or at least 100 times the number of genes to be tested to ensure that the empirical null distribution has resolution to calculate non-trivial p-values. An alternative approach we compare to is computing a null distribution for each gene. Based on numerical examples,the LS were insensitive to the distribution of the features after (see an example of LS from `GLISS` in Fig. 11).

**Figure 11:**
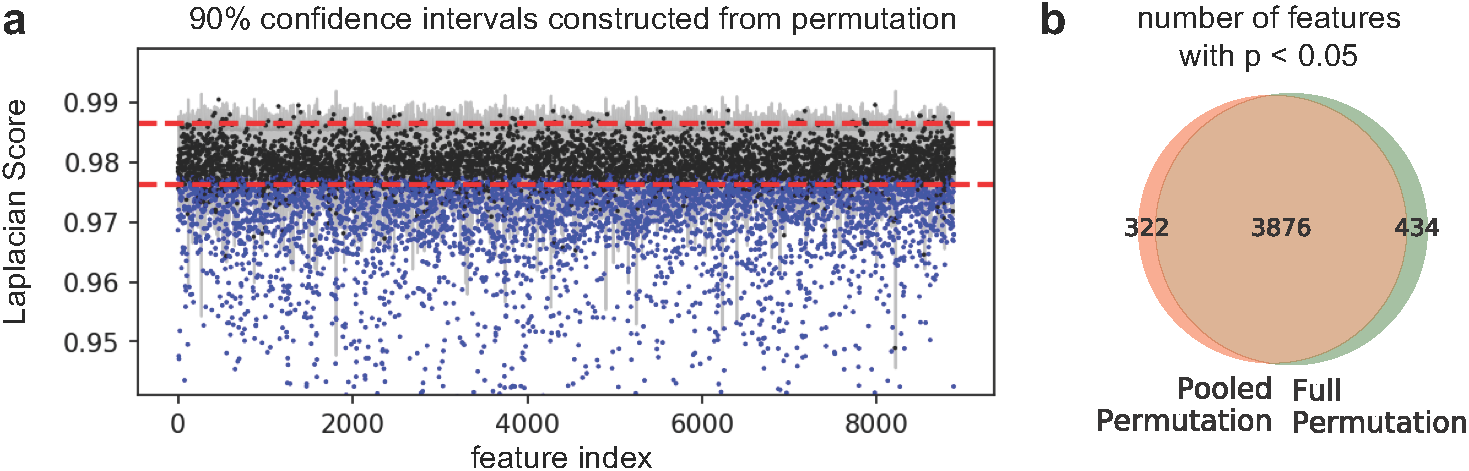
**(a)** We used the real data from [3] to compute the Laplacian Score between the initial graph 𝒢* and each of the 8,883 features, shown as individual data points. For the full simulation strategies, we permuted each feature 1,000 times to obtain feature-specific empirical null distribution. The 90% confidence for each feature (with 5% on each side) is shown by the gray bars. For the pooled simulation strategy, we randomly sampled 10,000 features to permute and computed a single empirical null distribution. The corresponding 90% confidence interval is highlighted by the red dashed lines (with 5% on each side). Note that the gray bars mostly agree with this interval. The features that have a full permutation p-value < 0.05 are indicated in blue, equivalent to the points that fall below their corresponding 90% interval. **(b)** Considering the features with p-value < 0.05, we find that those identified by the full permutation strategy highly overlap with those identified by the pooled permutation strategy.

## C Proof of Theorem 1

For the solution to Eq. (12) to be unique, G must have only one connected component, due to the following.

### Proposition C.1. (Number of connected components)

*Let 𝒢 be an undirected graph with non-negative weights. Then the multiplicity k of the eigenvalue 0 of* **L** *equals the number of connected components in 𝒢. Further, their corresponding eigenvectors span a k-dimensional subspace [26]*.

When there are *k* > 1 zero eigenvalues, the eigenspace of these k eigenvalues is spanned by the indicator vectors of those components: if *𝒜* is a set of connected nodes, an indicator eigenvector would be the vector with value one at the coordinates corresponding to nodes in 𝒜 and zero otherwise. However, any linear combinations of these indicator vectors could solve the first *k* problems in Eq. (10).

On the contrary, suppose now that there is only one connected component; then the second eigenvalue must be greater than zero. Because the stochasticity of the data generating process and the mapping from matrices to their spectra is continuous [47], all the eigenvalues will be distinct with probability 1. Next, we invoke the Spectral Theorem for Hermitian matrices as follows.

### Proposition C.2. (Unique eigenvectors)

*Let* **L** *be a symmetric matrix with distinct eigenvalues. Then its corresponding eigenvectors are orthonormal and are unique up to* ± *sign [48]*.

The uniqueness of the second eigenvector solved by Eq. (12) follows from this Spectral Theorem: since the sign change does not change the relative ordering of the data points, we have shown that **v**_2_ generates a unique ordering, which completes the uniqueness proof.

## D Details for Main Integrative Simulation

First, we specified a generative model that captures patterns found from the liver lobule study [3] for our simulation. Recall the generative model in Eq. (7), where *f*_*j*_ captures the dependency between the gene expression and the latent variable *ξ*. Because some of the features share the same *f*_*j*_, we use templates to describes these classes of features depend on *ξ* in the same way. Without loss of generality, we restrained *ξ* ∈ [0,1].

### Non-null templates

We set three monotonic templates for the LM genes (Figure 7A):

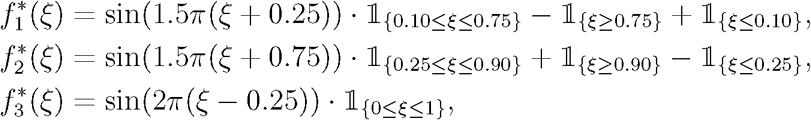

Here 𝟙_{𝒜}_ is the indicator function which equals to 1 if *𝒜* is true and zero otherwise. For the templates for the unknown non-null features (Figure 7A), we included six templates including those with more oscillating behavior. We had 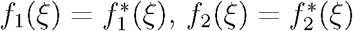, and 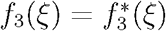 as above, and

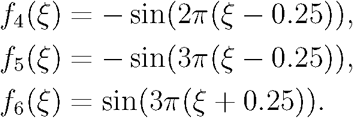

### Noise model

For the null variables, we used also used normal random variables. However, to include correlations within each 150-feature block, we generated samples from 𝒩 (0.5 · **1**_500_, Σ), where Σ equals to 1 on the diagonal and 0.5 otherwise. Regardless of sparsity of the non-null features, we kept the 150 block size for the null features to maintain the correlative effect. We also truncated each values to be between 0 and 1.

## E Details for Real Data Analysis

### E.1 The Hepatocyte Dataset

We downloaded the gene expression count matrix `table_s1_reform.txt` from the original publication [3]. As suggested by the authors, they had already filtered out the non-hepatocytes and performed moderate normalization to account for unwanted confounding effects. We processed the data using the `scanpy` Python package [29]. Using their built-in quality control procedure, we filtered out 16,998 genes that were expressed in less than 10 cells and performed log-transformation on the count data with ln(x + 1). We further filtered out genes that had standard deviation less than 0.1 in the log-transformed space. These filters led to 1,415 samples and 8,883 genes (which included the six landmark genes identified from [3]).

We used the default parameters for `GLISS` to analyze this gene expression matrix: we ran the SV gene selection step with the 6 predefined LM genes, and then we applied latent space inference and gene dimension reduction and clustering steps with the selected SV genes combined with the LM genes. To obtain the discrete spatial zones by [3], we downloaded their cell-zone probability matrix and used the mostly likely zone as the assignment. We did not perform any parameter tuning to analyze this dataset.

### E.2 The Enterocyte Dataset

Enterocytes constitute a large proportion of cells in the epithelial layer of the intestinal tract. The migration patterns of the enterocytes have been studied in the context of the villus axis: the position of the cell along the axis has been shown to correlate with multiple factors such as the age of the cells [2] (Figure 1B). To discover novel SV genes, Moor et al. characterized landmark genes with bulk RNA-sequencing of laser capture microdissected tissue (LCM-RNA-seq) [4]. Their main analysis linked their LCM-RNA-seq with scRNA-seq from enterocytes in a different study [43].

We first analyzed the SGE data, which consisted of bulk RNA-seq of 6 consecutive zones along the crypt-villus axis, with 3 replicates per zone, measuring a total of 20,000 genes. After quality control, approximately 1,000 were kept as candidates for LM genes, and `GLISS` selected over 300 LM genes from the SGE data. To identify the LM genes from SGE data for `GLISS`, we only considered genes with mean threshold greater than 4.5 after log-transforming the TPM values. Because there were 6 RNA-seq samples/zones and 3 replicates per sample, we only had 18 data points to construct a spatial graph. Thus, we created a Gaussian kernel graph instead of a k-NN graph, and selected the SV genes based on the default procedure with 10,000 permutations and FDR level 0.05. We identified 350 LM genes with which we used to analyze the scRNA-seq data for the remaining procedures under `GLISS`.

To reproduce the LM genes from the original study, we re-ran the original bulk RNA-seq analysis pipeline (as part of the MATLAB script under `matlab_03_zonation_reconstruction.m`, [4]) and obtained 109 landmark genes (out of 453 expressed genes) that consisted of 45 top landmark genes (tLM) and 65 bottom laandmark genes (bLM). tLMs were genes expressed in zone 1 with geometric mean < 2.5 and maximal mean expression above 10^−3^ after normalization; and tLMs were genes expressed in zone 5 with geometric mean > 3.5 and maximal mean expression above 10^−3^ after normalization. The corresponding enterocyte scRNA-seq dataset from [43] included 1,383 cells and 9,656 genes (obtained from the Seurat [13] processing pipeline described in [49]).

We obtained the processed scRNA-seq data, which included 1,383 cells and 9,656 genes, from the original study, which had already filtered unrelated cell types and normalized the expression values. For gene clustering, each gene was represented by 7 features, each corresponding to the average expression in a zone in the original study. Based on K-means clustering on the genes in this 7-D feature space, the workflow obtained 5 clusters of genes that share similar spatial patterns. To independently visualize the 7-D data, we used UMAP embeddings and discovered that the 5 gene clusters were separate but adjacent to one another. Using the spectral gap criteria, we initially identified 15 clusters (Extended Figure 16), but we chose the second largest gap of 9 clusters for comparison with the original study (Figure 10C).

## F Supplementary Figures

**Figure 12:**
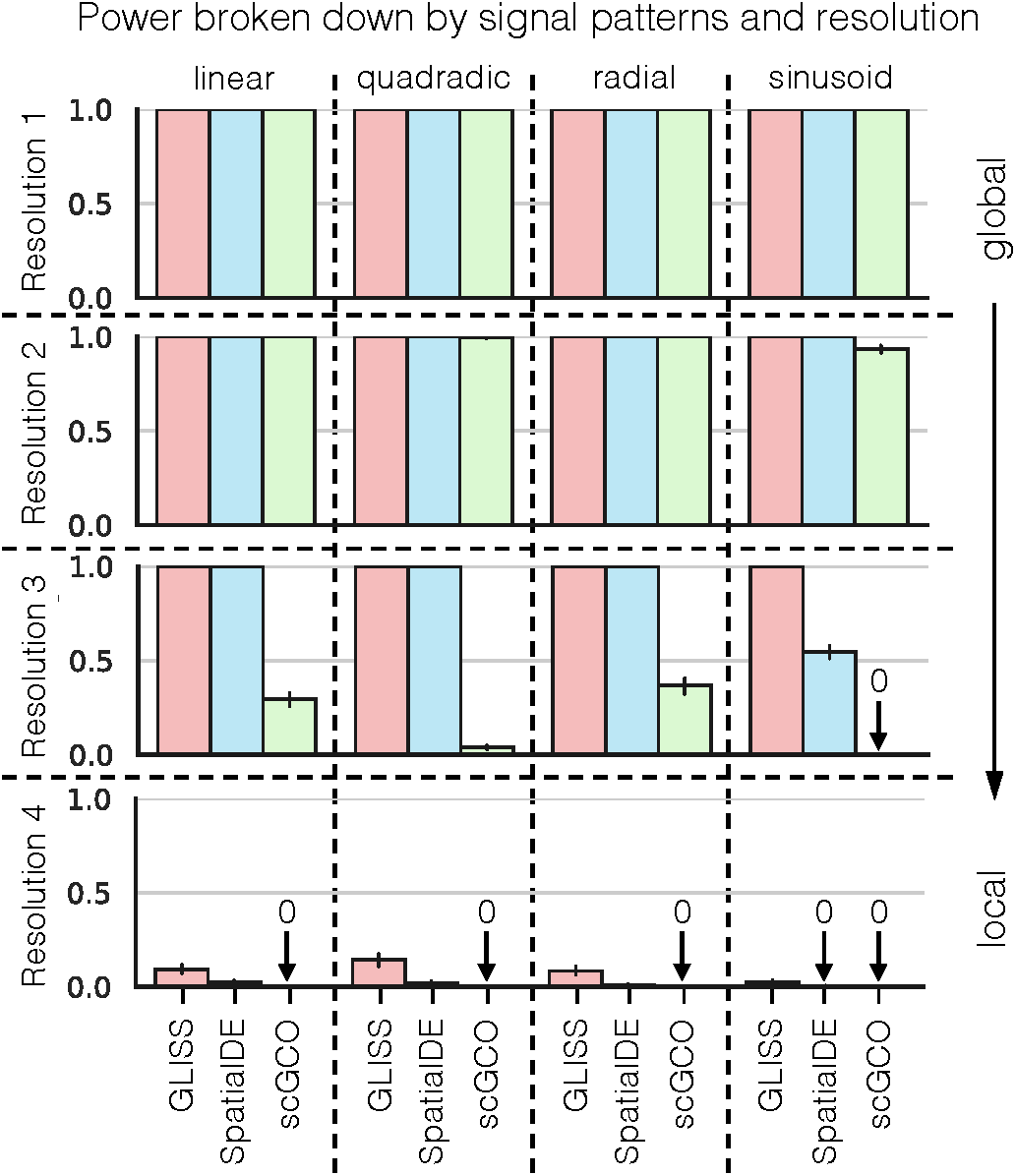
The empirical power broken down by different template resolutions. Each resolution consists of 25 genes, and the power is averaged over 20 trials with independent noise. Related to Figure 5

**Figure 13:**
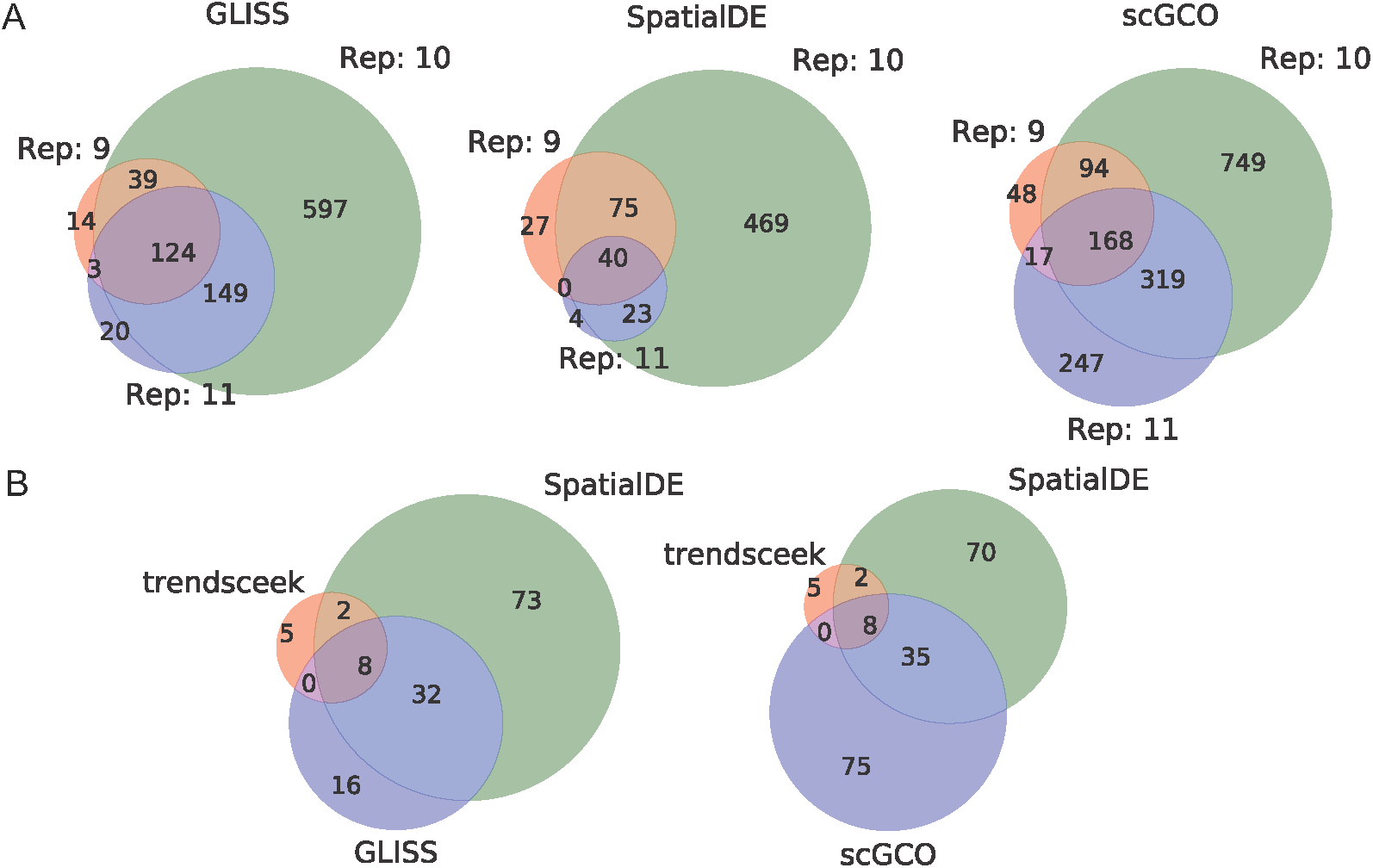
SGE data analysis with 3 real datasets. (A) The mouse olfactory bulb SGE dataset [15]. (B) The breast cancer biopsies SGE dataset [15]. Related to Figure 6.

**Figure 14:**
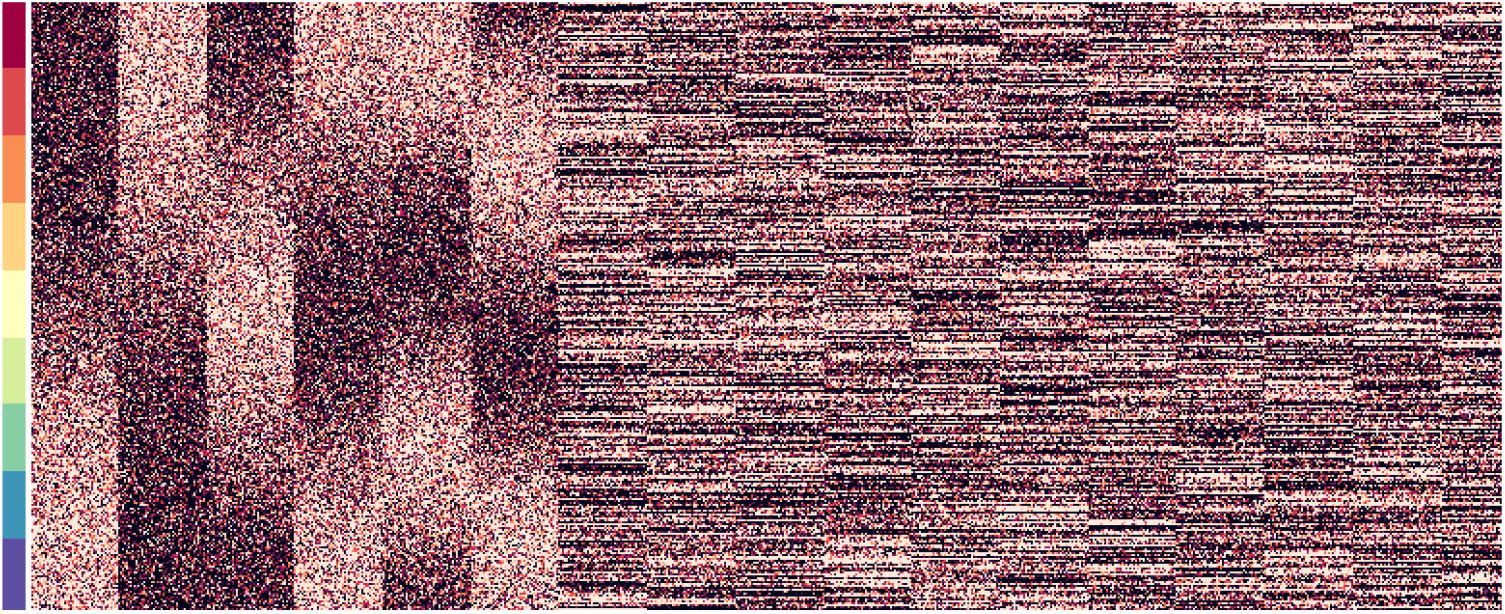
The simulated matrix consists of 1,500 samples (rows) and 6,000 features (columns). A (2,500-column) truncated heatmap is shown by including all the unknown non-null features organized by their templates, followed by the null features organized by their correlation structure. The colored axis indicates that the cells here are ordered by the position in the latent space. Related to Figure 7.

**Figure 15:**
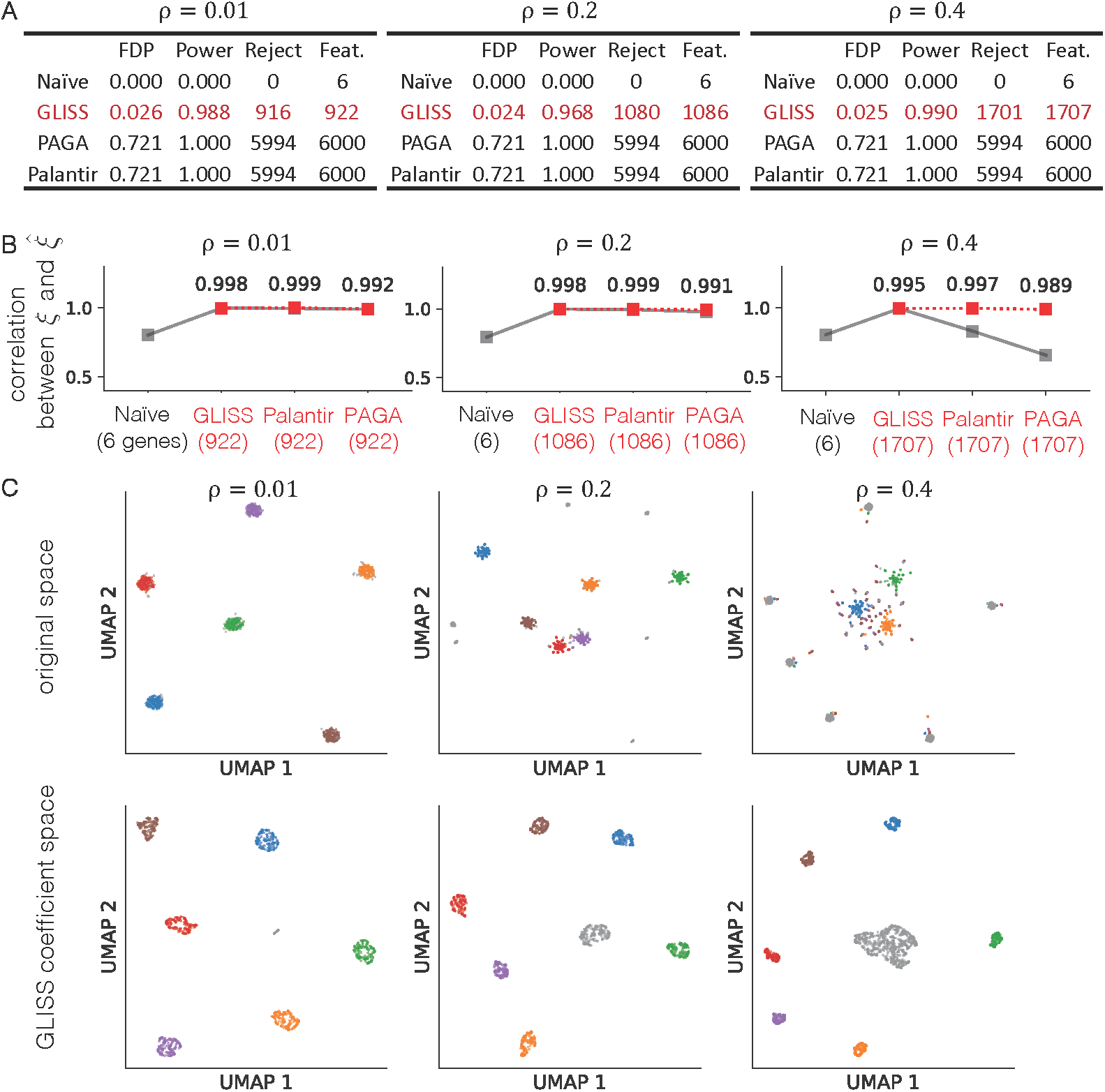
Additional simulation results for scRNA-seq. (A) Summary of the FDP, power, number of selected genes, and number of genes used for latent space inference. (B) The absolute Spearman’s correlation between the true *ξ* and the inferred variable 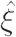. Here, `Palantir` and `PAGA` both take as input the expression of the LM and SV genes selected by `GLISS`. (C) The UMAP embedding of genes only the SV genes selected by `GLISS`. Feature selection made the non-null groups overall more distinguishable for the Naive approach (compared to Figure 7D). However, the Naive approach could not distinguish the six non-null gene clusters in the UMAP space as well as `GLISS` when *ρ =* 0.4. Related to Figure 7.

**Figure 16:**
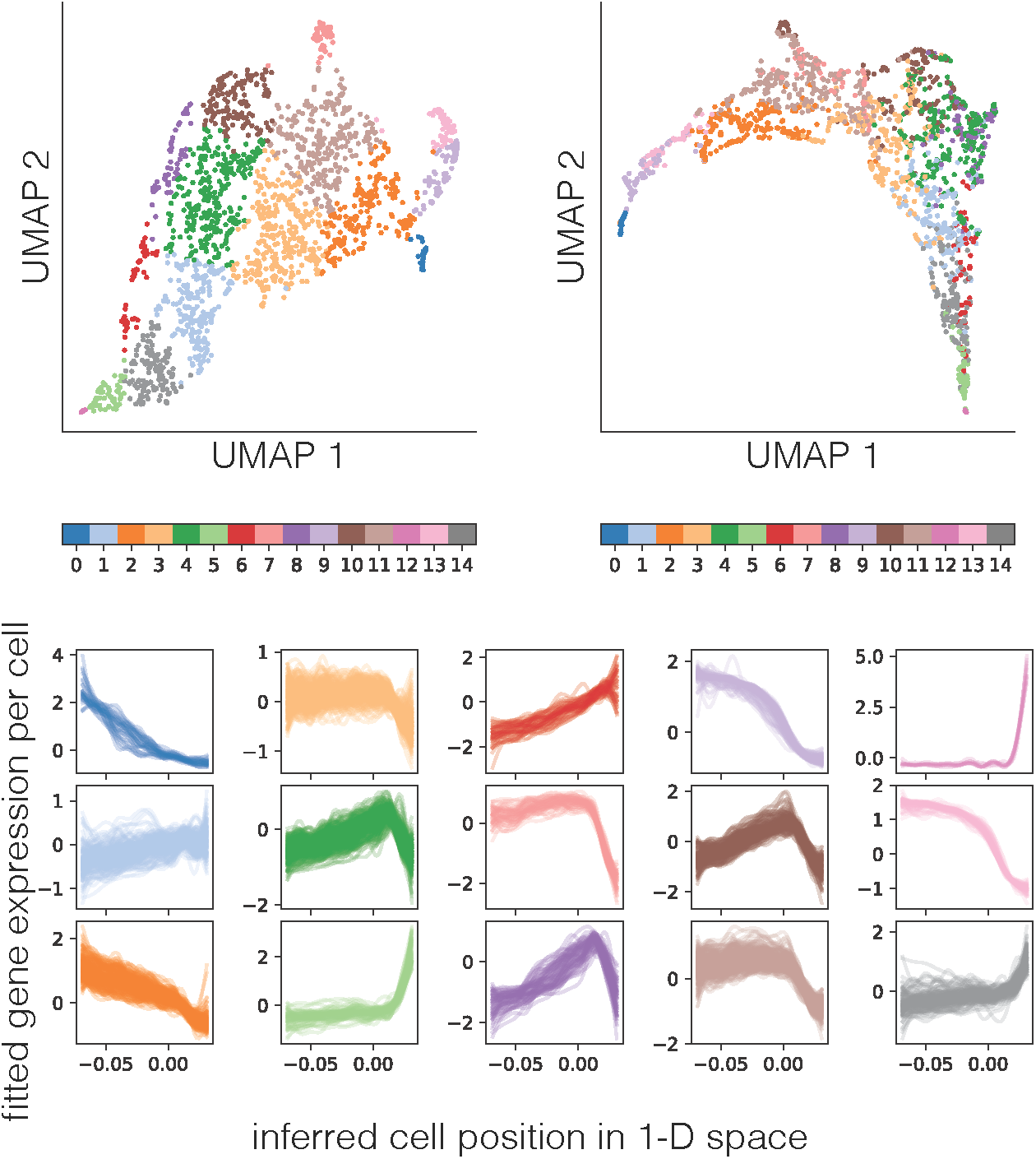
Gene dimension reduction result for the enterocytes with 15 clusters, which are more visible in coefficient space. Related to Figure 10.

## Footnotes

1 Although we use a nearest-neighbor graph like [26, 27], our approach applies to any distance-based graphs.

2 Under certain applications of SGE, each sample may correspond to the average expression of a group of cells in a region indexed by *i*, e.g. measured under bulk RNA-seq.

3 Often, we can rely on 3 or more LM genes to reconstruct interesting spatial structures.

4 One can substitute **Y** with the full gene expression matrix **X** prior to SV gene selection if needed, because each gene is fitted separately. Consequently, it would take more computation time if one decides to fit every gene rather than only the SV genes.

5 The FDP was computed accounting for null features that were correlated via noise: we count a null feature that is correlated with the LM genes as a true discovery.

## Notes

### Competing Interest Statement

The authors have declared no competing interest.

